# An integrated signaling threshold initiates IgG response towards virus-like immunogens

**DOI:** 10.1101/2024.01.28.577643

**Authors:** Wei-Yun Wholey, Alexander R. Meyer, Sekou-Tidiane Yoda, James L. Mueller, Raisa Mathenge, Bryce Chackerian, Julie Zikherman, Wei Cheng

**Affiliations:** Department of Pharmaceutical Sciences, 428 Church Street, University of Michigan, Ann Arbor, Michigan 48109, USA; Division of Rheumatology, Rosalind Russell and Ephraim P. Engleman Rheumatology Research Center, Department of Medicine, University of California, San Francisco, California 94143 USA; Department of Molecular Genetics and Microbiology, School of Medicine, University of New Mexico, Albuquerque, New Mexico 87131, USA; Department of Biological Chemistry, 1150 W. Medical Center Dr., University of Michigan Medical School, Ann Arbor, Michigan 48109, USA

**Keywords:** virus-like structures, epitope density, nucleic acid, antibody, Toll-like receptor, CD19, virus-like particles

## Abstract

Class-switched neutralizing antibody (nAb) production is rapidly induced upon many viral infections. However, due to the presence of multiple components in typical virions, the precise biochemical and biophysical signals from viral infections that initiate nAb responses remain inadequately defined. Using a reductionist system of synthetic virus-like structures (SVLS) containing minimal, highly purified biochemical components commonly found in enveloped viruses, here we show that a foreign protein on a virion-sized liposome can serve as a stand-alone danger signal to initiate class-switched nAb responses in the absence of cognate T cell help or Toll-like receptor signaling but requires CD19, the antigen (Ag) coreceptor on B cells. Introduction of internal nucleic acids (iNAs) obviates the need for CD19, lowers the epitope density (ED) required to elicit the Ab response and transforms these structures into highly potent immunogens that rival conventional virus-like particles in their ability to elicit strong Ag-specific IgG. As early as day 5 after immunization, structures harbouring iNAs and decorated with just a few molecules of surface Ag at doses as low as 100 ng induced all IgG subclasses of Ab known in mice and reproduced the IgG2a/2c restriction that has been long observed in live viral infections. These findings reveal a shared mechanism for nAb response upon viral infection. High ED is capable but not necessary for driving Ab secretion *in vivo*. Instead, even a few molecules of surface Ag, when combined with nucleic acids within these structures, can trigger strong antiviral IgG production. As a result, the signaling threshold for the induction of neutralizing IgG is set by dual signals originating from both ED on the surface and the presence of iNAs within viral particulate immunogens.

**One-sentence summary:** Reconstitution of minimal viral signals necessary to initiate antiviral IgG

## Introduction

Despite their enormous diversity, mammalian viruses have revealed surprising similarities in the host Ab responses for both the immediate and long-term following infection. In different animal species, there is a rapid induction of potent neutralizing IgG independent of CD4^+^ T cells^1–4^. In mice, this IgG is dominated by IgG2a/2c, the so-called ‘IgG2a/2c restriction’^5^ of murine Abs. The production of Abs is long-lasting after the resolution of primary infections^6–12^. Despite the fundamental importance of these phenomena to our understanding of host antiviral immunity and to the development of vaccines, the molecular determinants that explain these similarities have remained elusive. There are two intrinsic barriers that hinder our understanding of these similarities across different realms^13^ of viruses: (1) the diversity and sophistication of biological virions in their supramolecular structures^14^, and (2) the inherent complexity in studies of live viral infections, where inflammation^15,16^ and cytokine secretion^17^ can modulate the immune network to influence Ab responses.

To understand the common features in antiviral Ab responses, we hypothesize that these similarities may stem from B cell responses to the common biophysical and biochemical attributes shared by viruses across different realms, and these common attributes collectively convey an integrated signal at the single-cell level in Ag-specific B cells, which leads to potent antiviral IgG. In this study, we consider two common yet essential attributes of viruses: surface epitope density (ED, i.e., the average number of epitopes per virion) and internal nucleic acids (iNAs). As we know, all viruses display surface proteins with certain spatial density, which are required for viral entry to initiate infection of host cells; and all viruses contain nucleic acid (NA) genomes internal to the virions, which are required for replication of the viruses inside the host cell. In fact, this minimal consideration of common viral features is fully consistent with the recent formal definition of a ‘virus’ by the International Committee on Taxonomy of Viruses, in which viruses are defined operationally as “*a type of mobile genetic element that encodes at least one protein that is a major component of the virion encasing the nucleic acid of the respective mobile genetic element*…”^18^. Rigorous testing of our hypothesis requires examination of host Ab responses induced by ED and iNA, both independently and collectively, in a single system of virus-like immunogens, both at early time points and over the long term. Preferably, this test should be conducted for at least two different protein Ags and in hosts of different genetic backgrounds to support the broad relevance and validity of this hypothesis. Moreover, naturally occurring virions have densities of surface proteins that vary over several orders of magnitude,^19,20^ ranging from 200 to 30,000 molecules per µm^2^. The test of this hypothesis will thus involve systematic variation of ED over a range relevant to diverse virus structures.

There are additional immunological rationales that support both ED and iNA as potential determinants of antiviral Ab responses. First, it is well established that a high-density, ordered display of an epitope on virus-like particles can elicit strong Ab responses and even break Ab self-tolerance^21–28^. Ag valency, related to the concept of ED but not identical, has been shown to have profound effects on Ab responses^29–31^. In particular, we have showed recently that ED on liposomes alone, without any iNA, can trigger robust Ag-specific B cell signaling and proliferation *in vitro* by evading Lyn-dependent inhibitory pathways for B cell activation^32^. This mode of B cell activation by ED is fundamentally distinct from that of soluble proteins, which are - by contrast – constrained in a Lyn-dependent manner. These results indicate that ED alone, in the absence of iNA, can convey a unique signal of B cell activation not simply explained by B cell Ag receptor (BCR) crosslinking. Second, after the recognition of protein epitopes on virions by the BCRs and BCR-mediated endocytosis, the encapsulated NA inside virions can activate endosomal NA-sensing Toll-like Receptors (TLRs)^33^, which can produce a range of effects on B cells^34–38^. It has been established that innate sensing of microbial infections regulates adaptive immunity^39,40^. However, how these two common features of virions, ED and iNA, orchestrate antiviral Ab responses for both the immediate and long term to mediate host protection remains incompletely understood, in particular with regard to how BCR signaling triggered by varied ED is integrated with iNA-dependent signaling through endosomal receptors. The examination of ED in this context is especially relevant given the fact that naturally occurring virions have EDs that vary widely.^19,20^

Studies using live, attenuated or inactivated viruses have contributed substantially to our current understanding of Ab responses towards viruses^4,6,22,41,42^. However, it is difficult to isolate the roles of viral ED and iNA in these studies due to the complex integrated supramolecular structures of viruses. For example, bacteriophage Qβ-derived virus-like particles (VLPs) are highly valuable tools that have offered significant mechanistic understanding in Ab responses to virus-like immunogens, especially during the early phases of Ab responses^26,36,43,44^. However, Qβ VLPs are mixtures of phage proteins and single-stranded RNA (ssRNA) self-assembled from *Escherichia coli* cell culture in diverse forms^45^. Although these phage structural proteins are all foreign to mice or macaques, it has been challenging to assess the contribution of these phage structural proteins to Ab response in isolation due to the ability of phage coat proteins to bind ssRNA non-specifically. In fact, the ability to encapsidate foreign RNA is the basis on which the Qβ coat protein is used commercially for protection of heterologous RNAs^46^. Attempts have been made to digest away RNA packaged inside VLPs to assess the role of VLPs in the absence of NA^43^. However, the VLPs after digestion still elicited NA-specific IgG2a response in C57BL/6 mice^43^, and the basis for this NA-specific IgG2a remains unclear.

Moreover, Qβ is an icosahedral virus and it is known that the structural and mechanical features of icosahedral viruses are different from those of enveloped viruses^47^, in terms of the spatial density of epitopes^19^, rigidity and fluidity of the structures^48^, which are all likely to influence Ab responses^49^. These differences between icosahedral and enveloped viruses confer cautions in extending mechanistic insights obtained from Qβ-based VLPs to enveloped viruses, such as SARS-CoV-2, influenza viruses and HIV-1.

Existing vaccine platforms such as lumazine synthase^50^, ferritin nanoparticles^51^, or computationally designed self-assembling protein nanomaterials^52^ have also been extensively studied and used in vaccine design and development^53–58^. However, these structures lack iNA, which makes it difficult to assess the role of iNA in the context of virus-like immunogens using these structures.

To overcome these limitations, we have developed a system of synthetic virus-like structures (SVLS) based on liposomes to mimic the structural features of enveloped viruses that can be programmed in a modular fashion. These SVLS are constructed using highly purified biochemical ingredients and recapitulate the common biophysical features of naturally occurring enveloped viruses^59–61^. By contrast to bacteriophage Qβ-derived VLPs, the SVLS system uses lipid bilayers^28,62,63^ instead of phage structural proteins to serve as the scaffold for the display of Ags, eliminating Ab responses towards the scaffolds which will otherwise complicate data interpretation. The protein Ags of interest are then covalently conjugated onto the surface of these structures in an ordered array with a programmable density. To mimic naturally occurring viruses, we selected lipids that were abundant in natural plasma membranes to construct SVLS (Materials and Methods). We also avoided cationic lipids that are known to be immunostimulatory in nature^49^, which were extensively used recently for the therapeutic delivery of mRNA vaccines^64^. More importantly, this system allows us to encapsulate iNA with *all-natural* phosphodiester backbones inside SVLS, which mimics the spatial confinement and chemical composition of a typical viral genome^60,61^.

To investigate the individual as well as the collective roles of ED and iNA on host Ab responses, it is highly desirable to independently modulate both ED and iNA on these structures so that the resulting Ab response can be quantified in an animal model. Due to the modular nature of these structures, SVLS allow us to do so in a *single* system of virus-like immunogens. In this paper, we describe the early Ab responses initiated by SVLS as early as Day 5 (D5) after immunization. We present evidence that the signaling threshold for the induction of a rapid class-switched nAb response is set by dual signals originating from both ED on the surface and the presence of iNAs within viral particulate immunogens. In the companion paper, we describe and characterize the SVLS-induced Ab responses that have been followed for up to two years after immunization. These results reveal individual as well as collective contributions of essential viral features - titrated independently and across a broad range - towards antiviral Ab responses. The shared features in Ab responses as we will describe between two completely unrelated protein Ags indicate that our results are broadly applicable to understanding immune responses to viruses and vaccines.

## Results

### The characterization and the use of SVLS

We sought to generate reagents with highly purified components in order to model and modulate essential viral features in isolation and to co-ordinately define their impact on host Ab responses. To do so, we prepared four sets of SVLS for the current study: two of them, schematically shown in Fig. 1a and 1b insets and termed pRBD and pHEL, respectively, are SVLS that display either purified receptor-binding domain (RBD) of the SARS-CoV-2, the causative agent of the COVID-19 pandemic^65^; or a purified mutant version of hen egg lysozyme (HEL), a well-established protein Ag that has been used extensively in mechanistic studies^66^. RBD and HEL are not evolutionarily related, nor do they possess any significant sequence or structural similarities. Each protein was displayed in a specific orientation at programmed densities on the surface of SVLS. The interior of pRBD and pHEL was filled with phosphate buffered saline (PBS). The other two sets: pRBD(iNA) and pHEL(iNA), schematically shown in Fig. 1c and 1d insets, are SVLS that not only display RBD or HEL on their surface, but also encapsulate NAs inside their structures. All protein display was achieved through site-specific covalent conjugation on the lipid bilayer surface to ensure stability of the attachment with time^59–61^. No proteins other than RBD or HEL were present in these structures. Fig. 1a through 1d show dynamic light scattering measurements of these structures, with diameters around 120 nm, which is very close to that of HIV, influenza virus and SARS-CoV-2^67–69^. Fig. 1e shows a representative epi-fluorescence image of individual pRBD(iNA) stained by a fluorescent RBD-specific Fab (Extended Data Fig. 1). We determined the ED, i.e., average number of protein molecules per structure, using both ensemble and single-molecule fluorescence techniques that were established previously^70–72^.

**Fig. 1.**
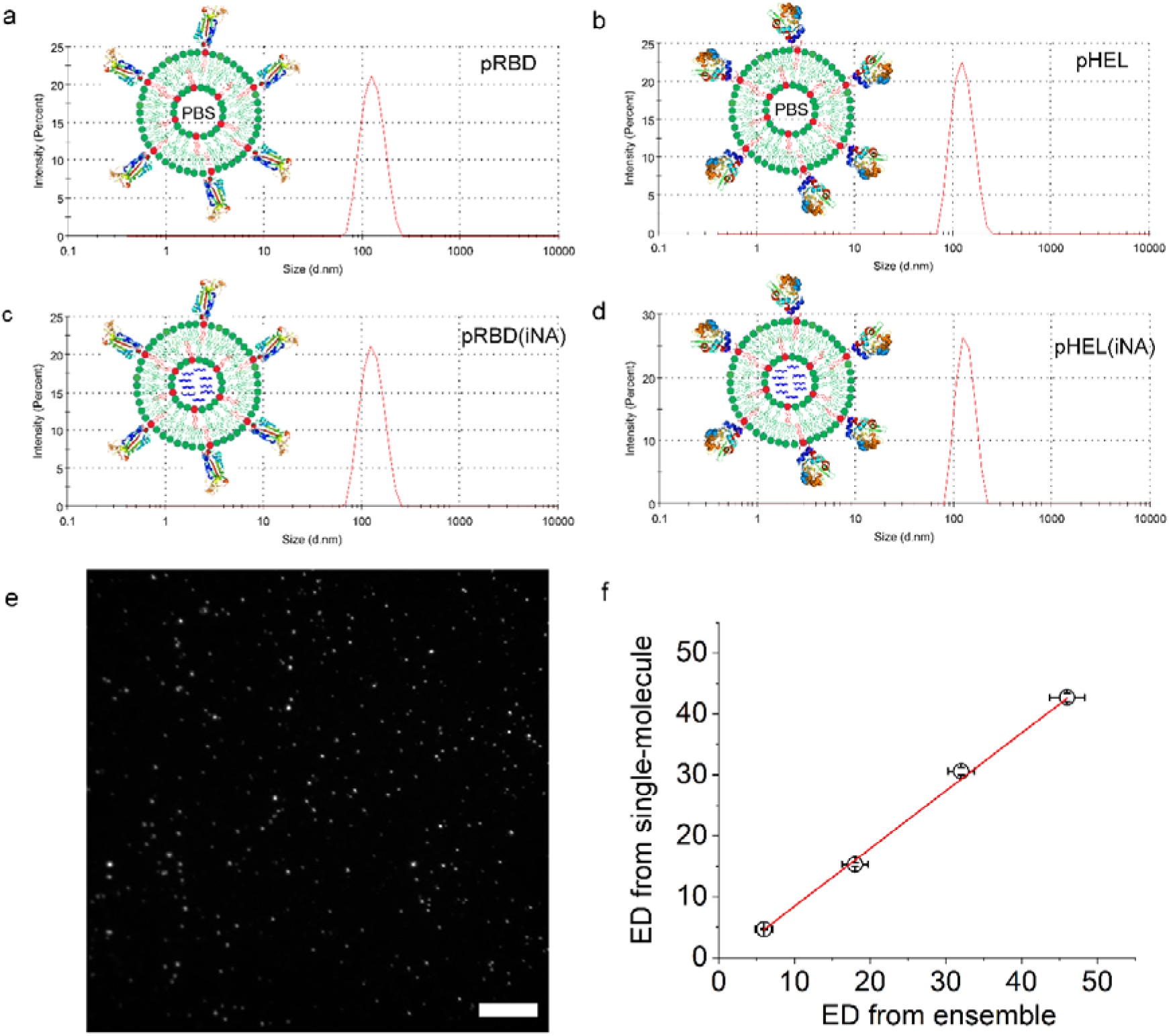
Characterization of SVLS that quantitatively incorporate the common consensus features of enveloped viruses. (**a**) through (**d**): representative intensity distributions of particle sizes measured using dynamic light scattering for four types of SVLS, schematically shown as insets in each panel. The structure surface is conjugated with RBD (a, c) or HEL (b, d) site-specifically at programmed spatial densities. The internal space of the structures is either PBS for investigation of ED in isolation (a, b) or loaded with nucleic acids to study the effects of iNA (c, d). The average diameters together with PDI values are 122 nm, 0.037 for pRBD; 120 nm, 0.039 for pHEL; 123 nm, 0.042 for pRBD(DNA1); and 124 nm, 0.054 for pHEL(DNA1). Cartoon insets adapted from ^60,61^. (**e**) A representative epi-fluorescence image of individual SVLS on a poly-L-lysine coated coverslip stained by a fluorescent Fab (Extended Data Fig. 1). The scale bar represents 10 µm. (**f**) Correlation between ED measured using single-molecule fluorescence technique (Extended Data Fig. 1) and the ensemble technique for a set of pRBD(DNA1). Error bars from respective measurements as described in Extended Data Fig. 1 and Table S1.

As shown in Table S1, we produced SVLS with a range of EDs for these structures that well encompass the EDs observed for naturally occurring viruses^19,20^. As shown in Fig. 1f, the values of ED for pRBD(iNA) that we measured using the ensemble-based method overall agreed with ED values from the single-molecule fluorescence method (Extended Data Fig. 1), with an adjusted R-square value of 0.997 from a linear regression analysis (the straight red line). The sensitivity of the single-molecule fluorescence method thus confirmed the SVLS with low ED values that we have prepared.

Most viruses infect their hosts through peripheral routes^47^. To simulate a natural viral infection, we delivered a submicrogram dose of Ag into mice through subcutaneous injection of SVLS. This allows viral collection by the initial lymphatics to encounter B cells in the lymph nodes^73^. A submicrogram dose of Ag is equivalent to picomoles of SVLS, which is a very low dose compared to those typically used for VLPs or other conventional model immunogens in mice^24,25,74,75^. Importantly, because natural viral infection does not involve any conventional adjuvants such as alum, throughout these studies, all the injections were performed without addition of any exogenous adjuvants. As a result, the injections of SVLS only introduced the highly purified protein Ags and lipids for pRBD or pHEL, and additional encapsulated NAs for pRBD(iNA) or pHEL(iNA).

### SVLS without iNA induce rapid IgG responses

In the absence of iNA or any other adjuvants, we found that a single injection of pRBD or pHEL at a submicrogram Ag dose is sufficient to rapidly elicit Ag-specific IgG responses in both B6 and BALB/c mice, provided that the ED on SVLS was above an apparent threshold^29^. This is shown in Fig. 2a and 2b for various conditions that we tested using pRBD and pHEL, respectively, in B6 mice. For both SVLS, the Ag-specific IgG could be detected as early as day 5 (D5) post injection, 45±6 ng/ml anti-RBD IgG (Fig. 2a Condition 6 circle) and 51±15 ng/ml anti-HEL IgG (Fig. 2b Condition 6 circle), well above the detection limit of ELISA (Extended Data Fig. 2). Because the reference monoclonal IgGs for ELISA are affinity matured, the true concentrations of Ag-specific IgGs are likely to be even higher than these values, which reflect rapid extra-follicular responses by naïve B cells, with stronger responses on Day 11 (D11) (Fig. 2a-2b, Conditions 6 and 7 columns). The Ag-specific IgG responses were absent in several controls including PBS (Condition 1), soluble proteins alone (sRBD or sHEL, Condition 2), or an admixture of soluble proteins and control SVLS without conjugated proteins on the surface or iNA (Condition 3). Increasing the Ag dose of either sRBD or sHEL by tenfold also failed to elicit detectable Ag-specific IgG, consistent with the known requirement for adjuvants to produce immune responses to soluble proteins in mice. The exquisite sensitivity of animals towards pRBD or pHEL *in vivo* are consistent with the distinct signaling mechanisms that we uncovered recently in the sensing of these virus-like immunogens by Ag-specific B cells *in vitro*^32^. In fact, our *in vitro* studies using pHEL revealed that these SVLS were over 1000-fold more potent than respective soluble proteins in the activation of Ag-specific B cells^32^. Fig. 2a and Fig. 2b also revealed apparent thresholds of ED for IgG induction by either pRBD or pHEL in B6 mice. At the same Ag dose, an ED less than 25 was not sufficient to induce an Ag-specific IgG above background for either pRBD or pHEL (Fig. 2a-2b, Conditions 4 and 5). Stronger IgG responses were elicited with increasing ED for both pRBD and pHEL above the respective thresholds.

**Fig. 2.**
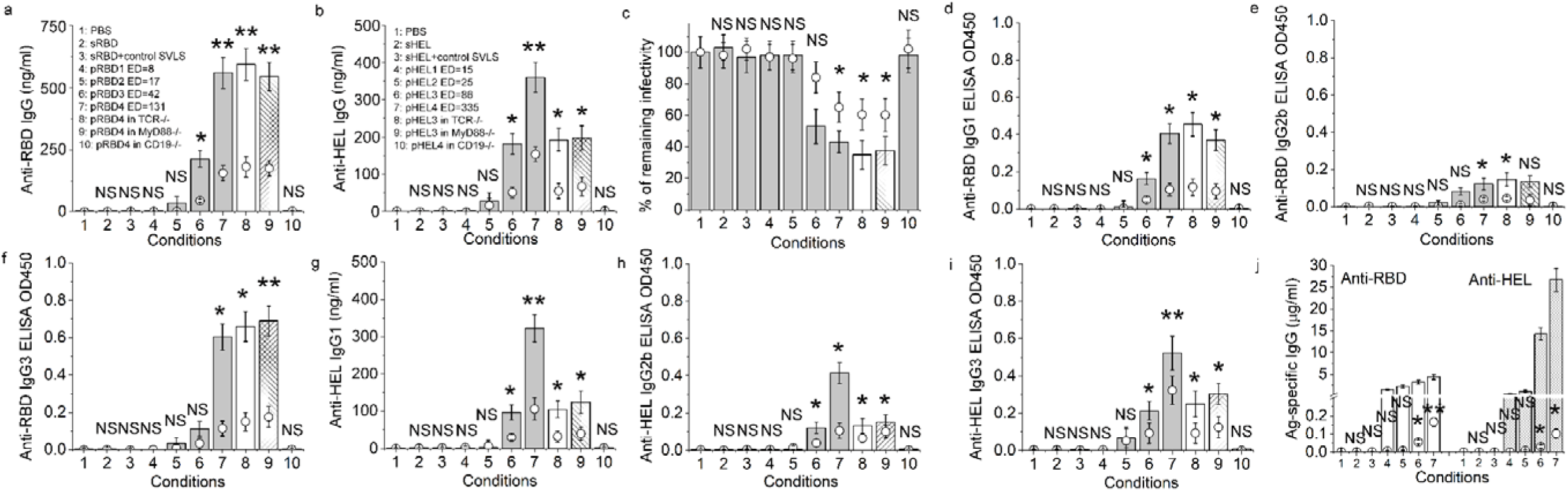
Rapid IgG responses induced by SVLS without iNA. (**a**)(**b**) RBD-specific IgG (a) or HEL-specific IgG (b) in mouse sera of B6 genetic background measured by ELISA after a single injection using various agents listed in the insets. (**c**) Neutralization of HIV-1 virions pseudotyped with SARS-CoV-2 S protein by mouse sera from various conditions in (a). (**d**) through (**i**): ELISA detection for RBD- and HEL-specific IgG subclasses in sera samples from (a) and (b). The y-axes for panels (d), (e), (f), (h) and (i) are shown in OD450 for 100-fold diluted sera because reference Ag-specific IgG subclasses are not available. (**j**) RBD-specific IgG (left) or HEL-specific IgG (right) in BALB/c mouse sera measured by ELISA after a single injection using various agents #1 through #7 as listed in (a) and (b) insets, respectively. Throughout Fig. 2, data from D5 post immunization are shown in circles and data from D11 are shown in columns; the doses of RBD and HEL were 0.24 and 0.3 µg per animal, respectively for Conditions 2 through 10. The pairwise statistical difference between each condition and Condition 1 on Day 5 was determined by Student’s T-test and shown using the following set of symbols; **: p-value < 0.01; *: p-value < 0.05; NS: not significant, p-value > 0.05. N=4.

In the context of viral infection, this immediate IgG secretion could offer numerous benefits to the host. Unlike IgM, the smaller size of IgG allows easy access to extravascular space and offers protection at those sites^76^. Class-switched B cells also have an advantage in signaling and survival over unswitched B cells due to the difference in the Ig cytoplasmic tail ^77^, which is critically important for signal amplification. Besides its role in direct neutralization of virion infectivity, IgG can hexamerize through its Fc after Ag binding, which leads to potent activation of the complement cascade^78^. It has been known for a long time that natural killer (NK) cells can bind to immune-complexed IgG via CD16-FcγRIIIA and this interaction activates the lytic mechanism of NK cells and also induces the production of lymphokines ^79,80^. Thus, the early secretion of IgG can produce multiple levels of effector functions for antiviral defense. Importantly, sera from mice immunized with pRBD (Conditions 6 and 7) could effectively neutralize HIV-1 pseudovirions^81^ that displayed the cognate S protein of the SARS-CoV-2 *in vitro* (Fig. 2c), suggesting that the serum Abs induced by SVLS would be functional *in vivo* to offer the host some protection against infection of SARS-CoV-2. For both pRBD and pHEL, this rapid response induced IgG1, IgG2b and IgG3 in B6 mice (Fig. 2d-i), but without IgG2c (Extended Data Fig. 3). This distribution is consistent with our previous report for a self-Ag displayed on SVLS in B6 mice^59^, where IgG2c is unique to immunization in the presence of NAs. Also, it shows similarity to the subclasses of IgG induced in the context of live viral infections^1^ yet distinct from those induced by the type II T-independent Ags^82^. These results indicate that in the absence of iNA, a foreign protein displayed on the surface of a virion-sized liposome can trigger the immediate production of neutralizing IgG *in vivo* to prepare for an antiviral defense, provided that the ED is above a threshold.

The presence of the HEL-specific IgG in B6 mice is intriguing because it is well established that the B6 strain is unresponsive to HEL due to its failure to generate a dominant MHC Class II determinant that is needed for recruitment of cognate T cell help^83^. The data thus suggest that the initiation of the fast IgG response by these SVLS may be independent of T cells. To test this formally, we immunized B6 mice that were deficient in both alpha/beta and gamma/delta T-cell receptors (TCR^-/-^)^84^ with either pRBD or pHEL. Statistical comparisons between Condition 7 (wild-type B6 or wtB6) and Condition 8 (TCR^-/-^) for pRBD4 in Fig. 2a or between Condition 6 (wtB6) and Condition 8 (TCR^-/-^) for pHEL3 in Fig. 2b for both D5 and D11 did not reveal significant differences (p-values>0.1 for all cases), suggesting that this IgG response was largely T-independent for both immunogens. Furthermore, Ab responses were also observed in B6 mice that were deficient in MyD88 (MyD88^-/-^)^85^ (Condition 9 in Fig. 2). Comparisons with respective wtB6 mice for both D5 and D11 did not reveal significant differences between Condition 7 (wtB6) and Condition 9 (MyD88^-/-^) for pRBD4 in Fig. 2a or between Condition 6 (wtB6) and Condition 9 (MyD88^-/-^) for pHEL3 in Fig. 2b (p-values>0.1 for all cases). These data ruled out the possibility that endotoxin or NA contamination of the SVLS produced these T-independent Ab responses and confirmed that the oriented display of surface epitopes above a threshold ED on a virion-sized liposome can provide a sufficient ‘*danger*’ signal to trigger rapid IgG secretion for antiviral defense, even though the magnitude of this response is not very high. Consistent with these results, our studies using pHEL on MD4 B cells revealed that pHEL alone, without any additional adjuvants, can trigger robust MD4 B cell proliferation and survival *in vivo*, and can do so *in vitro* in the absence of MyD88, IRAK1/4 and CD40 ligation^32^. Moreover, the Ab response induced by pRBD or pHEL was completely abrogated in B6 mice that were deficient in CD19 (CD19^-/-^)^86^ (Condition 10). CD19 is an important coreceptor on B cells^87^ that lowers the threshold for BCR activation^88,89^ and is essential for B cell responses to membrane-bound ligands^90^. These data indicate that CD19 is required for B cell production of IgG in response to a foreign protein displayed on the surface of a virion-sized liposome, even when the protein is displayed at a high density. Consistent with this, CD19^-/-^ MD4 B cell proliferation in response to pHEL was also highly impaired *in vivo*^32^.

Compared to wtB6 mice, BALB/c mice also mounted similar IgG responses on D5 after immunization with pRBD or pHEL (Fig. 2j circles). The IgG response included IgG1, IgG2b and IgG3, but without NA-specific IgG2a (Fig. 3). The presence of these similarities with B6 suggests that the early Ab response is an immediate host response that is conserved in both strains of mice. However, the IgG response in BALB/c mice on D11 was much higher than those in wtB6 mice (Fig. 2j columns), due to a much stronger IgG1 response that dominated on D11 for both pRBD and pHEL (Fig. 3a and 3e), suggesting that there are differences between the two mouse strains in the amplification of the Ab response beyond D5 after its initial activation.

**Fig. 3.**
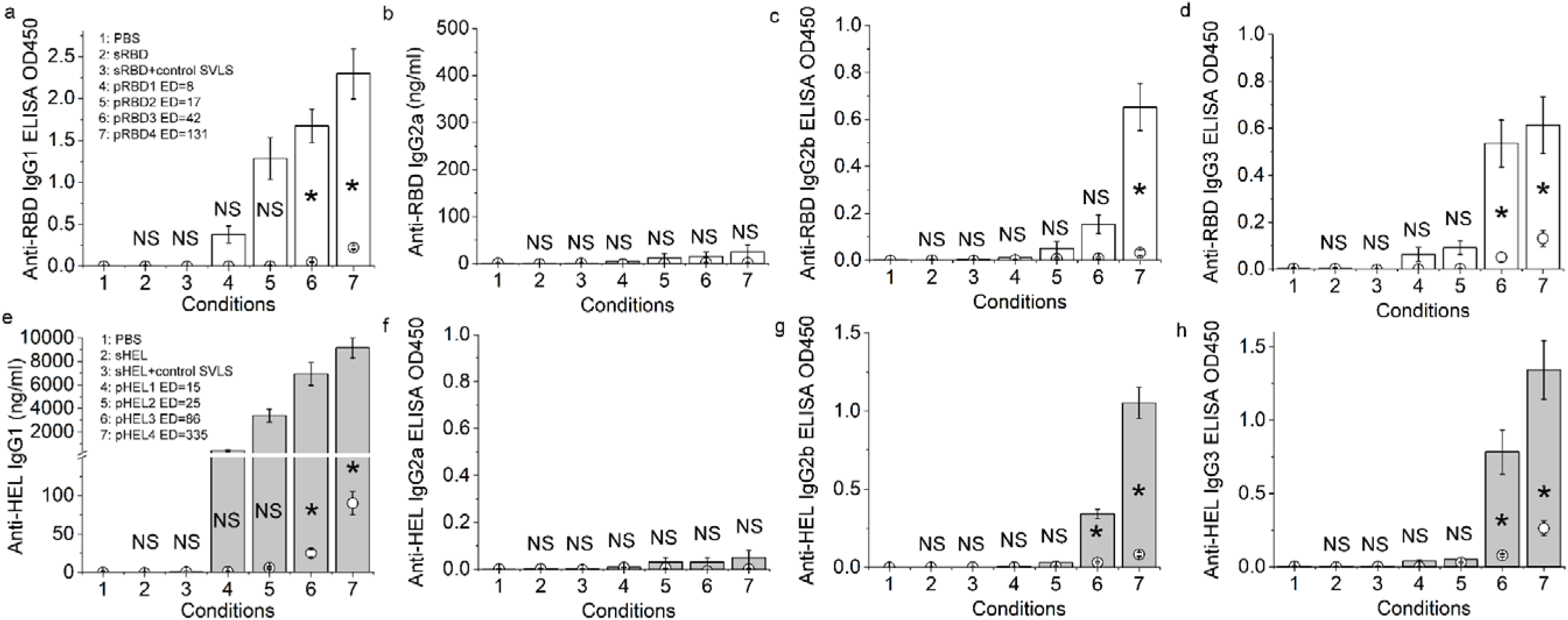
Ag-specific IgG subclasses in BALB/c mice upon a single subcutaneous injection of SVLS without iNA. (a) through (d) ELISA measurements for RBD-specific IgG1 (a), IgG2a (b), IgG2b (c), and IgG3 (d) in mouse sera after a single subcutaneous injection using various agents listed in (a) inset with no additional adjuvant. The dose of RBD per injection was 0.24 µg for conditions 2 through 7. (e) through (h) ELISA measurements for HEL-specific IgG1 (e), IgG2a (f), IgG2b (g) and IgG3 (h) in mouse sera after a single subcutaneous injection using various agents listed in (e) inset with no additional adjuvant. The dose of HEL per injection was 0.3 µg for conditions 2 through 7. Throughout Fig. 3, data from D5 post immunization are shown in circles and data from D11 are shown in columns. The pairwise statistical difference between each condition and Condition 1 on Day 5 was determined by Student’s T-test and shown using the same set of symbols as Fig. 2. N=4. The y-axes for all panels except (b) and (e) are shown in OD450 for 100-fold diluted sera because reference Ag-specific IgG subclasses are not available.

### SVLS with various iNA induce strong and rapid IgG responses

To examine the impact of iNA with natural phosphodiester backbones on the above IgG responses, we administered a single subcutaneous injection of pRBD(iNA) or pHEL(iNA) to mice at a submicrogram Ag dose without the use of additional adjuvants. We first measured the Ag-specific IgG responses in mice of B6 genetic backgrounds on both D5 (circles) and D11 (columns) post injection using ELISA, as shown in Fig. 4a and 4c for various conditions that we have tested for pRBD(iNA) and pHEL(iNA), respectively. These conditions included controls, SVLS of different ED, and SVLS containing different iNA sequences to model the diversity of viral genomes in either wtB6 or various gene knockout mice, as illustrated on the right in Fig. 4a and 4c. Many of these conditions induced strong Ag-specific IgG as early as D5 post injection. To quantify the amplification of the IgG response produced by iNA, we determined IgG titers for most conditions, as shown in Fig. 4b and 4d for pRBD(iNA) and pHEL(iNA), respectively. We observed a strong synergy between ED and iNA for SVLS encapsulating DNA1, a 20-mer single-stranded DNA that harbors two unmethylated CpG dinucleotides but has an all-natural phosphodiester backbone (Materials and Methods), as shown by Conditions 4 through 7 for both pRBD(DNA1) and pHEL(DNA1). When compared to either pRBD or pHEL without iNA, the titers of the IgG induced by these SVLS with DNA1 in wtB6 were all 10-to 1000-fold higher on D5 (circles, Fig. 4b and 4d), with even stronger responses on D11 (squares). As shown by several controls in Fig. 4a and 4c, the presence of iNA *internal* to SVLS was necessary for the strong and rapid responses observed, consistent with the requirement for B-cell intrinsic TLR activation. Condition 1 is an admixture of soluble proteins with DNA1. Condition 2 and 3 are admixtures of DNA1 and SVLS of different ED without iNA. Although unmethylated CpG DNA can activate TLR-9 ^91^, neither Condition 1 nor Condition 2 elicited Ag-specific IgG. Condition 3 elicited Ag-specific IgG but these responses were within error identical to those induced by SVLS alone without iNA (Fig. 2a and 2b), with p-values of 0.663 and 0.847 for pRBD3 on D5 and D11 and p-values of 0.728 and 0.787 on D5 and D11 for pHEL3 from one-way ANOVA tests. As we demonstrated previously^60,61^, when provided in a simple admixture, these NAs of all-natural phosphodiester backbones were quickly degraded in biological milieu, consistent with the apparent lack of their effects in Conditions 1, 2 and 3. In contrast, when encapsulated within SVLS, the lipid bilayer structures protected iNA from nuclease degradation^60,61^, similar to those in enveloped viruses. As a consequence, these NAs of all-natural phosphodiester backbones can unleash potent activity through intrinsic TLR activation after BCR mediated endocytosis of SVLS by the Ag-specific B cells *in vivo*.

**Fig. 4.**
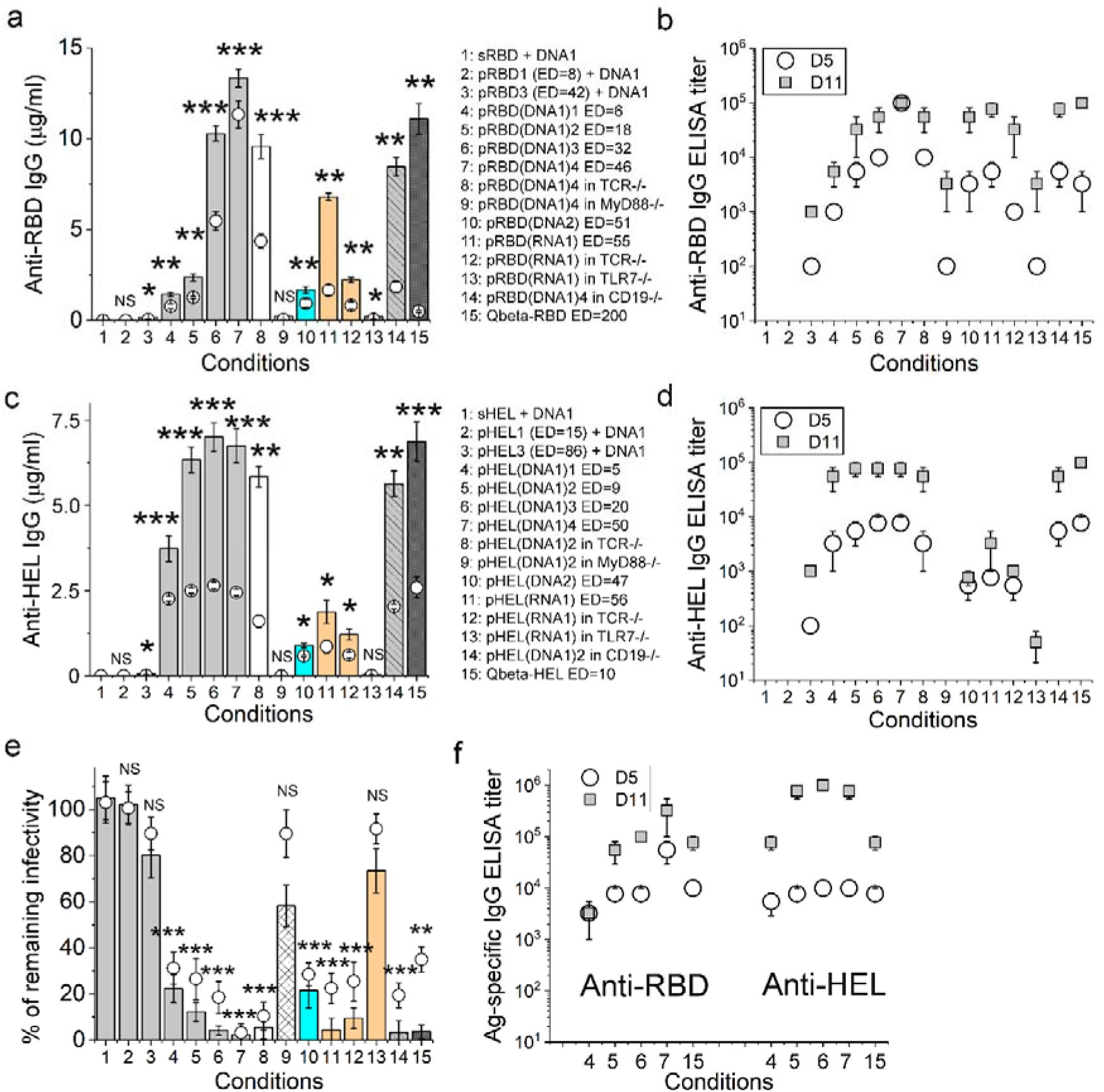
Strong and rapid IgG responses induced by SVLS with various iNA. (**a**) and (**c**) ELISA measurements for RBD-specific IgG (a) or HEL-specific IgG (c) in B6 genetic background after a single injection using various agents listed in respective insets. (**b**) and (**d**) Titers for RBD-specific IgG (b) or HEL-specific IgG (d) measured for various conditions as in (a) and (c). (**e**) Neutralization of HIV-1 virions pseudotyped with SARS-CoV-2 S protein by mouse sera from various conditions in (a). (**f**) Titer values for RBD-specific IgG (left) or HEL-specific IgG (right) in BALB/c mice after a single injection using agents #4, 5, 6, 7 and 15 as listed in (a) and (c) insets. Throughout Fig. 4, data from D5 post immunization are shown in circles and data from D11 are shown in columns or squares; the doses of RBD and HEL were 0.24 and 0.1 µg per animal, respectively for Conditions 1 through 15; the pairwise statistical difference between each condition and Condition 1 on Day 5 was determined by Student’s T-test and shown using the following set of symbols; ***: p-value < 0.001; **: p-value < 0.01; *: p-value < 0.05; NS: not significant, p-value > 0.05. N=4.

Naturally occurring viruses have diverse NA genomes. To test for broader relevance of the above results, we examined two other NA molecules: DNA2 and RNA1 (Fig. 4b and 4d Condition 10 and 11). DNA2 is a 20-mer single-stranded DNA that harbors two unmethylated GpC dinucleotides but has all-natural phosphodiester backbone, which was usually used as a control for CpG-containing DNA. RNA1 is a 20-mer ssRNA that is highly conserved among SARS-CoV-1, SARS-CoV-2 and MERS-CoV, and encodes motif V of an essential RNA helicase in these coronaviruses^92^ (Materials and Methods). Although the IgG response elicited by pRBD(DNA2) or pHEL(DNA2) was weaker than that of pRBD(DNA1) or pHEL(DNA1) at similar ED, DNA2 still enhanced IgG responses, indicating that the CpG dinucleotide motif was not required to observe this synergy. Specifically for pRBD(DNA2), the responses on both D5 and D11 were significantly higher than those induced by pRBD3, an SVLS of similar ED but without iNA (Fig. 2a Condition 6), with p-values of 0.0104 and 3.76e-4 from one-way ANOVA tests. For pHEL(DNA2), the responses on both D5 and D11 were also significantly higher than those induced by pHEL3, an SVLS of even higher ED but without iNA (Fig. 2b Condition 6), with p-values of 0.0494 and 0.0285 from one-way ANOVA tests. For pRBD(RNA1) (Fig. 4b Condition 11), although the IgG titer on D5 was lower than that induced by pRBD(DNA1)4, an SVLS of similar ED with DNA1 (Fig. 4b Condition 7, p-value=2.88e-8 from one-way ANOVA test), the IgG titer on D11 was statistically identical (p-value=0.356 from one-way ANOVA test). Therefore, CpG DNA is not unique in its strong potency for the synergistic induction of Ag-specific IgG responses. A natural viral genomic sequence is sufficient to induce a strong and rapid IgG response that is comparable in its magnitude.

The use of B6 mice allows us to dissect genetic requirements for the robust Ab responses induced by SVLS containing iNA [SVLS(iNA) herein]. First, the IgG response was largely T-cell independent, as shown by the strong Ag-specific IgG observed in T-deficient mice for pRBD(DNA1) and pRBD(RNA1) (Fig. 4b, Condition 8 and 12), pHEL(DNA1) and pHEL(RNA1) (Fig. 4d, Condition 8 and 12). In contrast, the IgG induction required MyD88 or TLRs, as shown by the much attenuated IgG response in mice deficient in MyD88 for pRBD(DNA1) or pHEL(DNA1) (Fig. 4a and 4c, Condition 9); and in mice deficient in TLR-7^93^ for pRBD(RNA1) or pHEL(RNA1) (Fig. 4a and 4c, Condition 13). However, in mice deficient in CD19, strikingly, the presence of DNA1 internal to these SVLS can elicit a fast and strong IgG response as early as D5 for both pRBD(DNA1) and pHEL(DNA1) (Fig. 4b and 4d, Condition 14, circles). On D11, the IgG titer induced by pRBD(DNA1)4 in CD19^-^/^-^ mice (Fig. 4b Condition 14) was as strong as that in wtB6 mice induced by the same agent at the same Ag dose (Fig. 4b Condition 7, p-value=0.356 from one-way ANOVA test).

Likewise, the IgG titer induced by pHEL(DNA1)2 in CD19^-^/^-^ mice on D11 (Fig. 4d Condition 14) was statistically identical to that in wtB6 mice induced by the same agent at the same Ag dose (Fig. 4d Condition 5, p-value=0.537 from one-way ANOVA test). These results are in sharp contrast to the requirement of CD19 by either pRBD or pHEL for induction of IgG in the absence of iNA (Fig. 2a-2b, Condition 10), and demonstrate the following: (1) a foreign protein on a virion-sized liposome alone requires CD19 to lower the BCR signaling threshold needed for IgG secretion. (2) The presence of iNA helps bypass this requirement for CD19 due to the stronger combined signals co-delivered by ED and iNA to an Ag-specific B cell. (3) At the level of individual B cells,^94^ BCR and intrinsic TLR activation feed into a common pathway at or before IgG secretion in response to virus-like immunogens *in vivo*. As a result, a deficiency in BCR-mediated signaling due to the absence of the BCR coreceptor CD19 (Condition 10, Fig. 2) can be compensated by the presence of iNA (Condition 14, Fig. 4b and 4d). In fact, we have showed recently that CD19 is dispensable for the immediate early (20 min) activation of MD4 follicular B cells in response to pHEL *in vitro*.^32^ This is also true when we probed these Ag-specific follicular B cells using pHELT in either heterozygous or homozygous context of CD19 deletion *in vitro* (Extended Data Fig. 4), in which pHELT displays a triple mutant of HEL (HELT) on surface with PBS in its interior. Altogether, these results show that CD19 is auxiliary for the activation of Ag-specific B cells in response to virus-like immunogens.

Supporting the convergence of signaling downstream of ED and iNA, we noticed that a very low ED of just a few molecules of Ag per SVLS was sufficient to elicit these strong IgG responses for both pRBD(DNA1) and pHEL(DNA1) (Fig. 4b and 4d, Condition 4). These results are surprising because an ED less than 25 was not sufficient to elicit an Ag-specific IgG above background for either pRBD or pHEL without iNA (Fig. 2). Consistent with these data, HEL-specific B cells are highly sensitive to ED but not affinity of the Ag display on pHEL or pHELT *in vitro* (Fig. 5). We can detect activation of HEL-specific B cells by pHEL or pHELT at ED’s as low as 3 molecules per structure *in vitro*, indicated by CD69 upregulation on these B cells by 4 hours after incubation with SVLS (Fig. 5). These data have several important implications regarding mechanisms of Ab responses to virus-like immunogens. First, the threshold of ED observed for IgG induction by SVLS without iNA (Fig. 2) is not equivalent to the threshold of ED required for BCR-mediated Ag endocytosis. Second, the threshold of ED for BCR-mediated endocytosis is low, because just a few molecules of Ag per SVLS can mediate the potent effects of iNA in SVLS. Third, the higher value of ED threshold required for IgG induction by SVLS without iNA likely reflects the magnitude of BCR-mediated signaling that is necessary to drive IgG secretion in the absence of TLR activation. Fourth, the amount of signal required for IgG secretion in Ag-specific B cells can be achieved with ED alone, at the expense of a relatively high ED value (Conditions 6 and 7 in Fig. 2), or the collective signals from ED and iNA together, which can work at a very low ED. These data are independent of the Ab response data in CD19^-/-^ mice as we just described above but support the same conclusion of convergent signaling between ED and iNA from a slightly different angle. Deficiency in BCR-mediated signaling, either due to low ED (Conditions 4 and 5, Fig. 2) or the absence of CD19, can be compensated by the presence of iNA. Together these data demonstrate that the signaling threshold for IgG induction is set by dual signals originating from both ED on the surface and the presence of iNAs within SVLS.

**Fig. 5.**
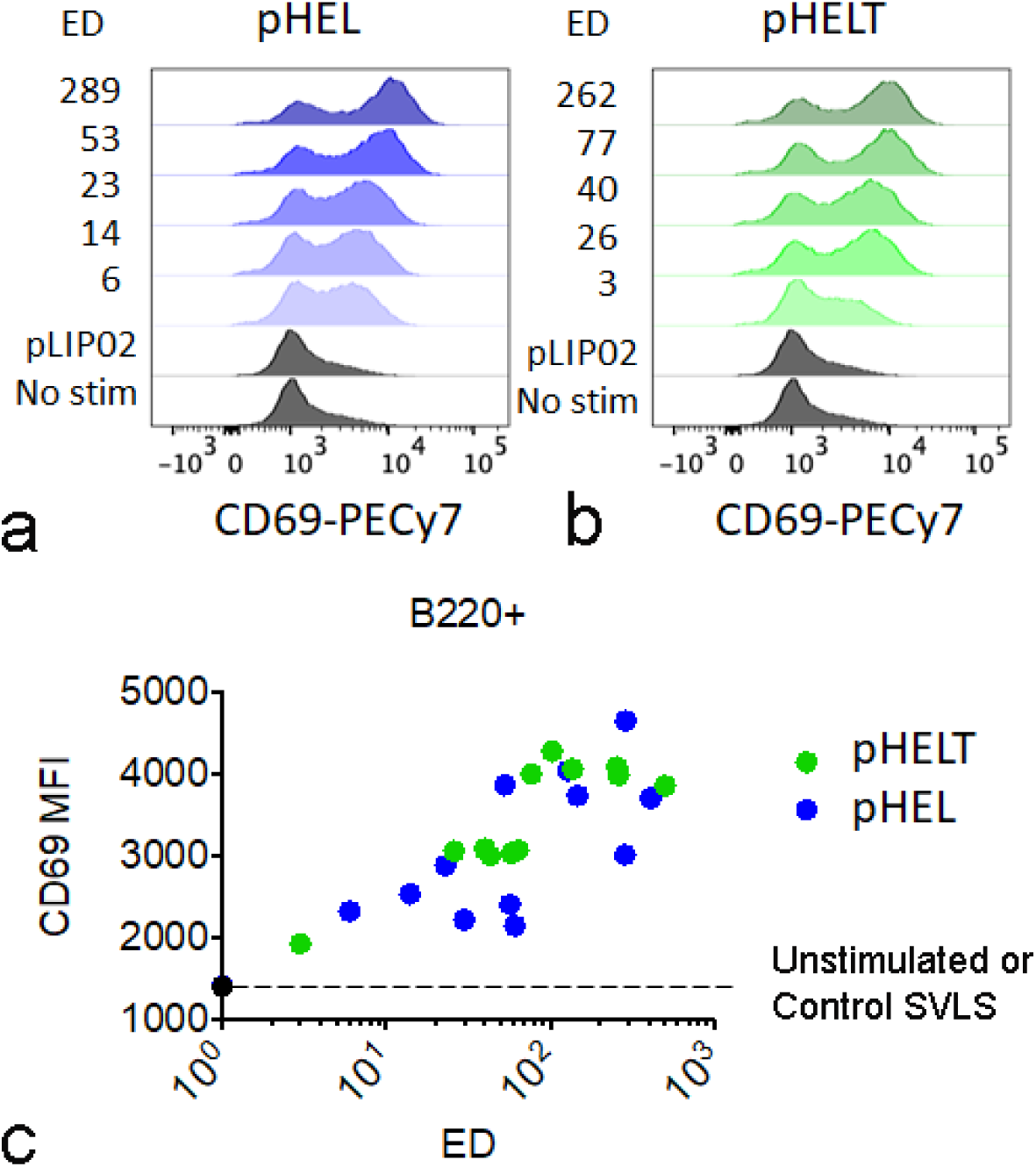
B cells are sensitive to ED but not affinity of Ag display on SVLS *in vitro*. Splenocytes and lymphocytes harvested from HyHEL10 mice (line harboring single copy of knock-in HC and Tg LC comprising Hy10 BCR that recognizes HEL) were incubated for 4 hrs with 1pM concentrations of pHEL (blue, epitope of intermediate affinity for the BCR) or pHELT (green, epitope of low affinity for the BCR) at varied ED’s. CD69 expression on B220^+^ live B cells was assessed by flow cytometry and is displayed in representative histograms, with panel **a** for pHEL and panel **b** for pHELT or quantified as mean fluorescence intensity (MFI, panel **c**). Data are representative of 3 replicates across 2 independent experiments.

As shown in Fig. 4e (Conditions 4-7, 10-11, the same set of conditions as in Fig. 4a), the strong IgG induced by pRBD(iNA) as early as D5 after injection can potently neutralize HIV-1 pseudovirions that display the cognate S protein of SARS-CoV-2 *in vitro*, with an even stronger potency on D11 after immunization. These strong neutralizing activities were obtained for all iNA sequences that we tested, suggesting a broad relevance of these results to understanding virus-mediated Ab responses. We noted that even in the absence of T cells, this IgG response was strong enough to potently neutralize cognate virions *in vitro* (Conditions 8 and 12 in Fig. 4e). These results are fully consistent with live viral infection literature showing potent neutralizing Ab (nAb) response in the absence of CD4^+^ T cells^1–4^. Along similar notes, protective nAb to influenza A virus can be produced in mice without T follicular helper cells^95^, and high-affinity nAb to SARS-CoV-2 can be made without T follicular helper cells in patients^96^. Furthermore, this fast and potent nAb response was also mounted in CD19^-/-^ mice (Condition 14 in Fig. 4e), demonstrating that the BCR coreceptor CD19 can be bypassed for an immediate and yet potent nAb response.

How does the Ab response induced by SVLS compare to other VLPs? We chose the commonly used bacteriophage Qβ VLPs platform^26,36,43,44^ to answer this question. The strong IgG induced by SVLS(iNA) was comparable in titers to those elicited by RBD or HEL conjugated on bacteriophage Qβ VLPs at the same Ag dose on both D5 and D11, despite the complete absence of the bacteriophage Qβ coat proteins in these SVLS(iNA). This was shown by Condition 15 in Fig. 4b for a Qβ-RBD conjugate with an ED of 200 and Condition 15 in Fig. 4d for a Qβ-HEL conjugate with an ED of 10 in wtB6. One-way ANOVA tests on these titer values revealed that Qβ-RBD induced the same level of RBD-specific IgG titers on D5 as pRBD(DNA1)1 and pRBD(DNA1)2, which were lower than those for pRBD(DNA1)3 and pRBD(DNA1)4; and Qβ-RBD induced the same level of RBD-specific IgG titers on D11 as pRBD(DNA1)3 and pRBD(DNA1)4, which were higher than those for pRBD(DNA1)1 and pRBD(DNA1)2 (Table S2a). For HEL, Qβ-HEL induced the same level of HEL-specific IgG titers on D5 and D11 as all pHEL(DNA1) (Table S2b). These results suggest that the SVLS platform has captured essential ingredients in conventional VLPs for potent IgG induction. It is feasible to elicit IgG responses of a similar magnitude to bacteriophage VLPs using SVLS(iNA), despite the fact that the epitopes in these two platforms have very different two-dimensional mobilities.

Like we observed in the B6 background, strong and fast IgG responses were also induced by SVLS(iNA) in BALB/c mice as early as D5. As shown in Fig. 4f, BALB/c mice mounted strong IgG responses towards pRBD(DNA1) of varied ED (Fig. 4f left, Conditions 4-7) with comparable magnitudes to B6 on both D5 and D11. Likely due to the available T cell help in BALB/c mice for HEL, the IgG responses towards pHEL(DNA1) in BALB/c mice were even stronger on D11 than those in wtB6 (Fig. 4f right, Conditions 5-7), indicated by one-way ANOVA tests on these titer values (Table S3). Therefore, an IgG response was robustly triggered upon a single exposure to a submicrogram dose of Ag on SVLS(iNA) in the absence of any other adjuvants in both strains of mice. The similar magnitudes of these responses between B6 and BALB/c mice suggest that this rapid and strong IgG secretion in response to the combined signals from ED and iNA in SVLS is conserved between the two strains and is highly efficacious against pseudovirions. Moreover, for both RBD and HEL, these responses induced by SVLS(iNA) could be quantitatively compared to those induced by either Qβ-RBD (Fig. 4f left, Condition 15) or Qβ-HEL conjugate (Fig. 4f right, Condition 15) at the same Ag dose in BALB/c mice, confirming our results for this comparison in wtB6 mice. Collectively, these data reveal that the SVLS platform can rival conventional VLPs for potent IgG induction.

Different IgG subclasses can mediate specialized effector functions^97^ that are important for antiviral defense. Both pRBD(iNA) and pHEL(iNA) induced IgG1, IgG2b, IgG2c and IgG3 in wtB6, with IgG2c being the dominant subclass for almost all the SVLS(iNA) (Fig. 6a through 6h). Analogous to wtB6, the IgG response in BALB/c induced IgG1, IgG2a, IgG2b, and IgG3, with IgG2a being the dominant subclass (Fig. 7). These results are fully consistent with the literature that IgG2a/2c is the predominant antiviral IgG in mouse upon infection by different viruses^5^. It is notable that SVLS(iNA) can induce all the IgG subclasses known in mice and reproduce this dominant pattern of IgG2a/2c, the so-called ‘IgG2a/2c restriction’^5^ of murine Abs observed in varieties of live viral infections, in our context without live infection nor any other adjuvants. All iNAs, including DNA1, DNA2 and RNA1, induced IgG2c in B6 genetic background (Fig. 6c and 6g), which was in sharp contrast to the undetectable IgG2c upon pRBD or pHEL immunization without iNA (Extended Data Fig. 3a and 3b). Therefore IgG2a/2c is uniquely induced via innate sensing of iNA^43^. Furthermore, incorporation of iNA lowers the ED thresholds for each IgG subclass. With iNA, the level of IgG1 is now comparable between wtB6 and BALB/c particularly for pRBD(DNA1) on both D5 and D11 (Table S4 for one-way ANOVA tests). Moreover, the comparison between SVLS(iNA) and Qβ-conjugated Ags reveals that Qβ was a stronger inducer of IgG1 after D5 of immunization in both wtB6 (Fig. 6a and 6e, Condition 15) and BALB/c mice (Fig. 7a and 7e, Condition 6). One possibility for this difference is the presence of multiple copies of the bacteriophage coat protein in Qβ VLPs, a foreign protein that is known to recruit T cell help in mice^59^.

**Fig. 6.**
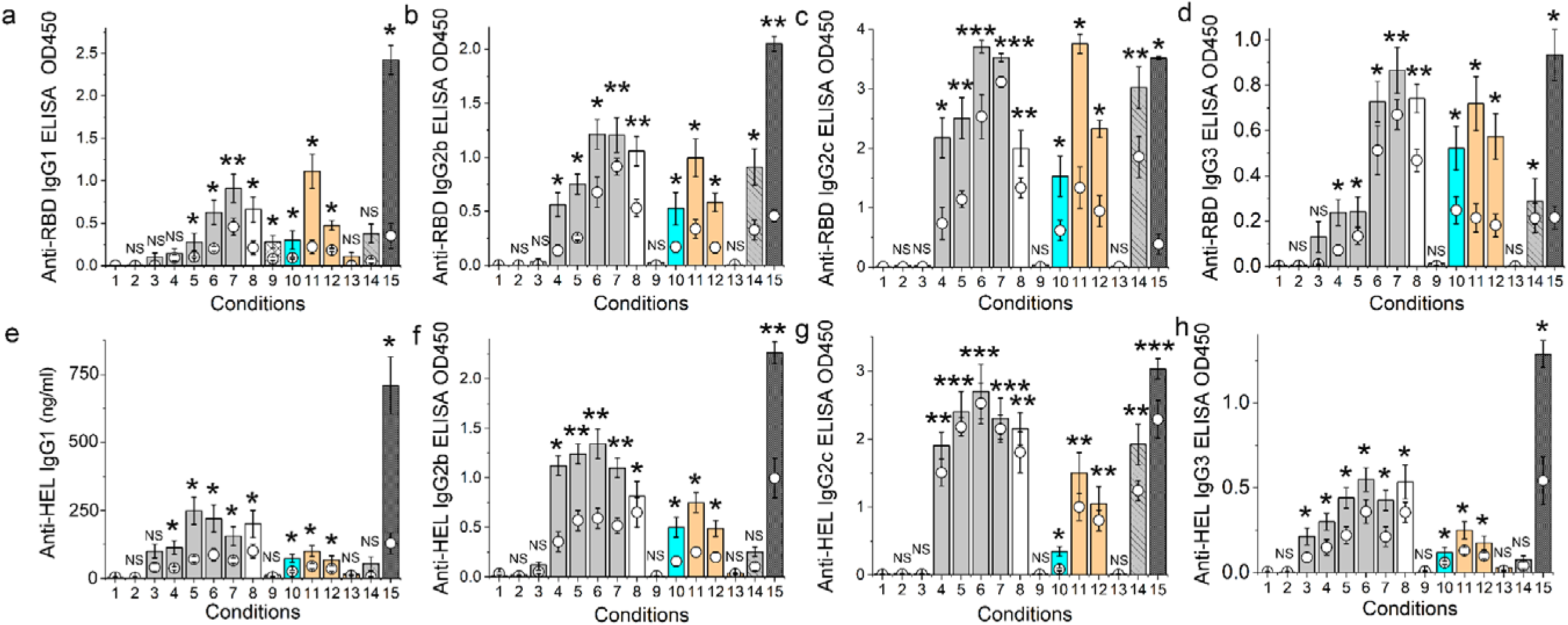
Ag-specific IgG subclasses induced by SVLS with various iNA in mice of B6 background. ELISA detection for RBD-specific (a through d) and HEL-specific (e through h) IgG subclasses for sera from conditions listed in Fig. 4a and 4c, respectively. The y-axes for all panels except (e) are shown in OD450 for 100-fold diluted sera because reference Ag-specific IgG subclasses are not available. Throughout Fig. 6, data from D5 post immunization are shown in circles and data from D11 are shown in columns. The pairwise statistical difference between each condition and Condition 1 on Day 5 was determined by Student’s T-test and shown using the following set of symbols; ***: p-value < 0.001; **: p-value < 0.01; *: p-value < 0.05; NS: not significant, p-value > 0.05. N=4.

**Fig. 7.**
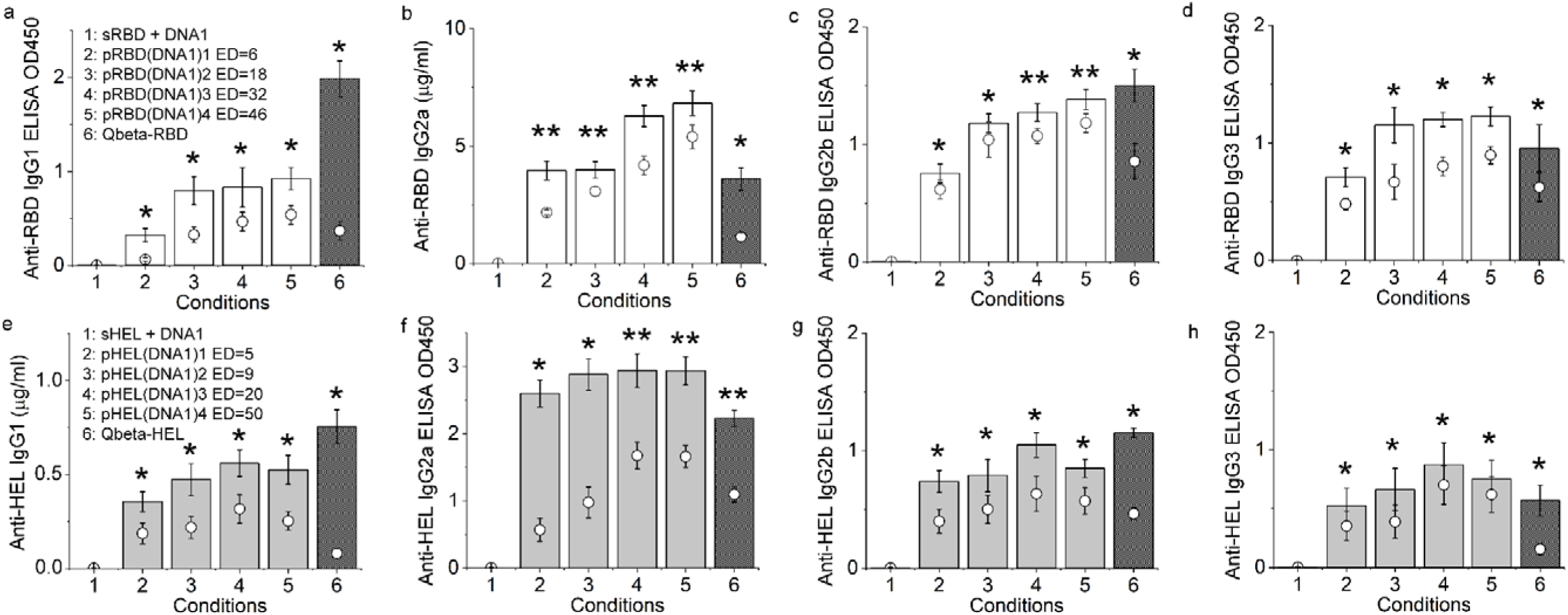
Ag-specific IgG subclasses in BALB/c mice upon a single subcutaneous injection of pRBD(DNA1), pHEL(DNA1) or Qβ conjugates. (**a**) through (**d**) ELISA measurements for RBD-specific IgG1 (a), IgG2a (b), IgG2b (c) and IgG3 (d) antibody in mouse sera after a single subcutaneous injection using various agents listed in (a) inset with no additional adjuvant. The dose of RBD per injection was 0.24 µg for conditions 1 through 6. (**e**) through (**h**) ELISA measurements for HEL-specific IgG1 (e), IgG2a (f), IgG2b (g) and IgG3 (h) antibody in mouse sera after a single subcutaneous injection using various agents listed in (e) inset with no additional adjuvant. The dose of HEL per injection was 0.1 µg for conditions 1 through 6. The y-axes for all panels except (b) and (e) are shown in OD450 for 100-fold diluted sera because reference Ag-specific IgG subclasses are not available. Throughout Fig. 7, data from D5 post immunization are shown in circles and data from D11 are shown in columns. The pairwise statistical difference between each condition and Condition 1 on Day 5 was determined by Student’s T-test and shown using the same set of symbols as Fig. 2. Error bars represent the standard errors (N=4). The dose of DNA1 in the admixture with soluble proteins was 1 nmol.

## Discussion

Viral infections of hosts produce Abs with shared features. Although these features have long been recognized in the literature^1–4,6,8^, the molecular determinants that can explain these shared features have remained elusive. Using a minimal and highly purified system of SVLS that enables us to independently vary ED and iNA, the two common attributes of all virions, we have characterized the individual as well as the collective effects of ED and iNA on host Ab responses. This manner of dissection for minimal determinants of Ab responses towards virus-like immunogens has not been possible before without the use of SVLS. Our data here demonstrate that the shared features in host Ab responses can be explained by B cell responses to the two common attributes of virion particles: ED and iNA in the context of intact virus-sized structures, without the need of active viral replication. This study also uncovered several new mechanistic understandings of Ab responses to virions, as we describe below.

### (1) Virus-like immunogens induce IgG *in vivo* independent of T cells or TLR activation

In response to viral infection, a fast T-independent IgG secretion could offer numerous benefits to the host, including direct neutralization of virion targets, easier access to extravascular space than IgM to offer protection^76^, better signaling and survival of Ag-specific B cells^77^ and direct activation of NK cells^79,80^ to mount an antiviral defense. Studies of Ab responses upon viral infection have revealed that for the majority of viruses, there is a rapid induction of potent neutralizing IgG independent of CD4^+^ T cells^1–4^ within just a few days after infection^42^. However, the mechanisms responsible for these IgG responses have been less clear. It was concluded that factors induced during live virus infection appeared to be required to induce the IgG^4^. In a recent study, it was shown that bacteriophage Qβ VLPs could induce IgG in the absence of T cell help^98^. However, because Qβ VLPs possess both high ED and ssRNA packaged inside the particles, which could serve as a potent TLR-7 ligand, it remained unclear if ED alone could induce IgG in the absence of T cells *in vivo*. Here using SVLS, we showed that antiviral IgG responses can be induced by a foreign protein displayed on the surface of a virion-sized liposome in the absence of T cells or TLR engagement, provided that the ED is above a threshold. This is true for both RBD and HEL, two unrelated Ags, suggesting that this may be a general phenomenon for Ab responses to Ags displayed on virus-sized structures. In fact, these results are also consistent with our prior report, where a liposomal display of a self-Ag can induce an IgG response^59^, although the Ag dose for the self-Ag was much higher than the Ag dose for the two foreign proteins in the current study. In the absence of iNA, the oriented display of proteins above a threshold ED on a virion-sized liposome can induce IgG1, IgG2b and IgG3 in both wtB6 and BALB/c mice, true for both foreign Ags in this study and also the self-Ag that was reported previously^59^. Thus, neither live infection nor any other adjuvants are necessary to initiate this IgG secretion. Together, these results demonstrate that the oriented display of viral surface proteins above a threshold ED on a virion-sized structure is sufficient to serve as a stand-alone ‘*danger*’ signal to trigger an IgG response in the absence of T cells or TLR signaling.

Consistent with this view and as we showed recently, MD4 B cells use a signaling mechanism that is distinct from that for sHEL to sense pHEL, notably through evasion of Lyn-dependent inhibitory pathways to initiate proliferation^32^. Furthermore, pHEL without iNA could maximally activate the NF-κB signaling pathway in MD4 B cells in a manner that mimics T cell help independently of MyD88, IRAK1/4 and CD40 ligation^32^. We hypothesize that the NF-κB signaling triggered by ED in the absence of iNA can drive the expression of activation-induced cytidine deaminase, the enzyme required for class-switch recombination (CSR)^76,99,100^, which then initiates CSR and IgG secretion *in vivo*. Alternatively, cytokine production *in vivo* by B cells or other immune cells may contribute. These possibilities remain to be tested.

### (2) Role of CD19 in Ab responses towards virus-like immunogens

CD19 is an important coreceptor for BCR activation^87,88,90^. It lowers the threshold of signaling for BCR activation by complement-coated Ag^88^, which is essential for B cell activation in response to membrane-bound Ags *in vitro*^90^ and crucial for Ab response to T-cell-dependent Ags *in vivo*^87^. However, the role of CD19 in host sensing and response to virus-like immunogens has not been established. Here we show that CD19 is required for Ab responses to SVLS *in vivo* in the absence of iNA (Fig. 2) but can be bypassed in the presence of iNA (Fig. 4). Consistent with these results, adoptive transfer studies using purified MD4 follicular B cells showed the requirement of CD19 for sustained B cell proliferation *in vivo* in response to pHEL^32^. At a mechanistic level, these results imply that CD19 helps amplify ED/BCR-mediated signaling, which is required for IgG secretion in the absence of iNA. However, in the presence of strong TLR-activating iNA, this CD19-mediated amplification of BCR signaling becomes redundant and thus dispensable. In particular, our finding that SVLS(iNA) can induce potent IgG response without CD19 (Fig. 4) may be useful for patients who suffer from Ab-deficiency syndrome due to mutations in the *CD19* gene^101^. These patients responded poorly to vaccines such as rabies^101^. In this case, an alternative vaccine formulation strategy to include potent TLR-activating iNA in SVLS may be useful for generating a better Ab response.

### (3) The signaling threshold set by dual signals from ED and iNA to initiate IgG responses

The synergy between a protein Ag and a TLR agonist in a B cell Ab response has been well recognized in literature ^35–38,102,103^. On a similar note, recently work by the Casali group also showed that BCR-TLR4 co-engagement can induce CSR and hypermutated nAb response in the absence of T cells^104^. However, our data showed that iNA can compensate for the deficit in ED/BCR-mediated signaling to allow for the potent induction of IgG secretion, which was not known previously. These results in both wtB6 and CD19^-/-^ mice have several implications regarding the initial events in Ag-specific B cell activation. First, BCR-mediated endocytosis can occur with as low as a few molecules of Ag per virion-sized structure *in vivo*. However, this low ED alone is insufficient to trigger IgG secretion, which requires a much higher threshold of ED, as schematically shown in Fig. 8. This result pertains to both basic B cell biology and also antiviral Ab responses. Studies have attempted to resolve the minimum BCR signaling threshold required to internalize Ags *in vivo*^105^. One study has suggested the presence of such a threshold in the avidity of BCR-Ag interactions^35^. However, this question has not been resolved due to the intrinsic complexity in B cell biology. Hou *et al*. reported that BCR signaling and internalization are mutually exclusive events, i.e., each BCR can undergo only one of two events, signaling or internalization, but not both^106^. This suggests that the threshold for BCR-mediated internalization may not be equivalent to the threshold for BCR signaling. Consistent with this view, our results here clearly showed that *in vivo*, the threshold of ED required for BCR-mediated internalization is much lower than the threshold of ED alone required for IgG secretion.

**Fig. 8.**
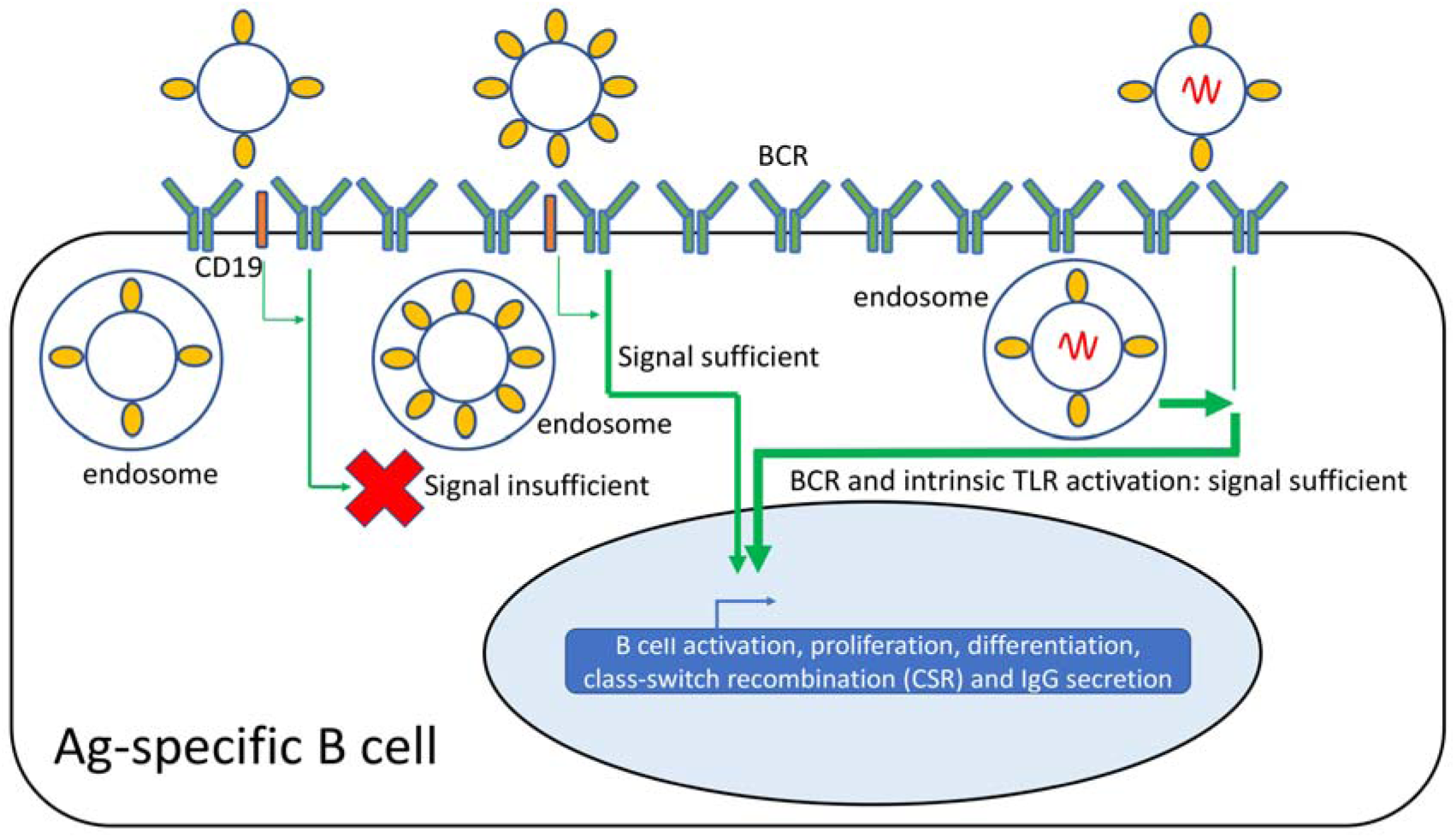
Illustration of signaling threshold required for IgG secretion in Ag-specific B cells upon recognition of virus-like immunogens *in vivo*. BCR-mediated endocytosis can occur to all particles depicted above, from low to high ED. However, for SVLS without iNA, both CD19 (slim orange rectangles) and a threshold of ED are required to meet the signaling threshold required for CSR and IgG secretion. For SVLS with iNA, CD19 can be bypassed for CSR and IgG secretion because the combined signal from BCR and intrinsic TLR activation can be sufficient to overcome the signaling threshold. Note that BCR and intrinsic TLR activation converge at or before CSR and IgG secretion take place. In response to virus-like immunogens, CSR and IgG secretion can occur independently of cognate T-cell help or TLR activation. Also, Ag-specific B cells may undergo proliferation and/or differentiation before CSR and IgG secretion, which is noted above.

Second, the capability of iNA to compensate for the deficiency of ED to mediate IgG secretion indicates that at a single-cell level, ED and iNA-mediated signaling converge to a common pathway that eventually leads to IgG secretion (Fig. 8). This quantitative coordination between ED and iNA has several implications for our understanding of B cell responses to virus-like immunogens. Naturally occurring virions have densities of surface proteins that vary over several orders of magnitude, ranging from 200 to 30,000 molecules per µm^2^ ^19,20^ and yet little is known about how this varied spatial density of viral surface Ags contributes to the immunogenicity of viruses. The capability of mouse B cells to respond to virus-like immunogens with as few as several molecules of surface Ag indicates that they can respond to a broad spectrum of viruses regardless of the densities of viral surface Ags, provided that there are potent TLR-activating iNA sequences inside these virions that are also accessible for TLR activation. On the other hand, because this mechanism is so potent in nAb induction, viruses that are known to co-evolve with their hosts may evolve to optimize their genome sequences to minimize potent TLR-activating iNA sequences such as CpG dinucleotides, or to encode virulence factors to antagonize this mechanism. In fact, it has been found that CpG dinucleotide sequences are suppressed in the genomes of virtually all eukaryotic viruses with a genome size < 30 kb^107^, including that of HIV-1^108^. Moreover, viruses may evolve to control the accessibility of their genomes for recognition by TLRs, given the fact that ligand availability for TLR activation is finely regulated *in vivo* to balance outcomes in antiviral responses and autoimmunity^109^. One major difference between SVLS and natural viruses is the accessibility of iNA. In SVLS, the iNA is accessible for recognition by TLRs upon BCR-mediated endocytosis. In natural viruses, many viral genomes are packaged in association with viral structural proteins. While the host animal may encode molecules^110^ or use other cells^111^ to regulate this important process of antiviral Ab production, one strategy that viruses can evolve is to preclude the access of viral genomes by TLRs until the virion is ready to uncoat its genome to start its life cycle of replication. Given the huge differences in the magnitudes of Ag-specific IgG with and without iNA, the modulation of TLR activation may naturally give rise to distinct levels of antiviral Ab responses. Whether this mechanism works in naturally occurring viruses remains to be tested.

## MATERIALS AND METHODS

### Synthesis of SVLS

All SVLS used throughout this study were prepared following protocols we published previously ^59–61^. These SVLS were constructed using nonimmunogenic lipids, with phosphatidylcholine and cholesterol comprising ≥ 99% of all lipids. These structures display protein antigens (Ags) of choice in a specific orientation on their surfaces with a regulated ED that quantitatively mimics the surface of naturally occurring viruses ^19,20^ (Fig. 1). The HEL recombinant proteins used in this study are (1) HEL, which contains two site-directed mutations at R73E and D101R, and (2) HELT, which contains three site-directed mutations at R21Q, R73E, D101R. Both were overexpressed in *E. coli* and purified to >95% purity following our established protocols ^60^. The SARS-CoV-2 RBD recombinant protein used in this study was overexpressed using 293F cells and also purified to >95% purity following our established protocols ^61^. DNA1, DNA2 and RNA1 were custom synthesized by IDT. The sequence of DNA1 is as follows, 5’-TCCATGA<u>CG</u>TTCCTGA<u>CG</u>TT-3’. The sequence of DNA2 is as follows, 5’-TCCATGA<u>GC</u>TTCCTGA<u>GC</u>TT-3’. The sequence of RNA1 is as follows, 5’-ACUGUUGAUUCAUCACAGGG-3’. Epitope density (ED) was quantified using methods that we established previously ^59–61,70^ that were also validated by single-molecule fluorescence techniques (Extended Data Fig. 1) that were developed and established in house ^71,72^. For SVLS with iNA, the average number of iNA molecules per SVLS was also quantified using methods that we established previously ^60,61^. Control SVLS had no surface protein conjugation nor internal encapsulated agents.

### Conjugation of RBD or HEL to bacteriophage Qβ virus-like particles (VLPs)

The bacteriophage Qβ VLPs were prepared as described ^112^. The purified RBD or HEL recombinant proteins, which each contained a single reactive cysteine near their respective C-termini, were conjugated to the surface of the VLPs using the heterobifunctional crosslinker Succinimidyl-6-[(β-maleimidopropionamido)hexanoate] (SMPH). Briefly, the VLPs were first derivatized with SMPH at a 10-fold molar excess of SMPH over Qβ coat protein. The mixture was incubated at 22°C for two hours and the excess crosslinker was removed by centrifugation at 4°C in an Amicon Ultra-4 centrifugal unit with a 100 kD cutoff. The RBD or HEL protein was then added to the derivatized VLPs and the mixture was incubated at 4°C overnight for RBD or at 20°C for 1 hour for HEL. Free proteins were then removed by centrifugation at 4°C in an Amicon Ultra-4 centrifugal unit with a 100 kD cutoff. The quantity of conjugated proteins was assessed using a denaturing polyacrylamide gel based on the intensity from silver staining in comparison with a standard curve obtained from the same gel.

### Mice

All animal procedures for mouse immunizations were approved by the University of Michigan Animal Care and Use Committee. Female C57BL/6 or BALBc/J mice (8 weeks, Jackson Laboratory) were used for immunizations. Prior to inoculation, all injection samples were filtered through a 0.45 µm pore size membrane to eliminate potential microbial contamination. 100 µL samples at a dose between 0.1 and 0.48 µg of respective protein Ags were injected into each mouse subcutaneously, 50 µL in each flank. Throughout the entire study, no adjuvants other than the SVLS were administered unless otherwise noted and only a single injection was administered throughout. Mouse blood was collected submentally using Microvette serum collection tubes (Sarstedt) three days before the first injection, and 5 days thereafter following specified blood collection schedules. The sera were harvested by centrifugation at 10,000 g for 5 minutes, aliquoted and immediately frozen and stored at - 80°C.

Four strains of gene knockout mice were purchased from Jackson Laboratory: Tcrb^tm1Mom^/ Tcrd^tm1Mom^ (#002122) (TCR^-/-^ herein) was deficient in both alpha beta and gamma delta T-cell receptors ^84^. Myd88^tm^^1.Defr^ (#009088) (MyD88^-/-^ herein) encodes a deletion of exon 3 of the myeloid differentiation primary response gene 88 ^85^. Cd19^tm1^(Cre)^Cgn^ (#006785) (CD19^-/-^ herein) has a Cre recombinase gene inserted into the first exon of the CD19 gene thus abolishing CD19 gene function ^86^. Tlr7^tm1Flv^ (#008380) (TLR7^-/-^ herein) has a segment of exon 3 of the Toll-like receptor 7 gene replaced by a lacZ gene and a loxP-flanked neomycin resistance cassette thus abolishing the function of Toll-like receptor 7 ^93^. Gene knockout mice were housed in a germ-free environment and immunization protocols were carried out as described above.

For B cell activation experiments *in vitro*, IgHEL (MD4) mice were previously described ^113^. HyHEL10 knock-in mice (Hc/+ knock-in and Lc+ Tg+) mice were previously described and generously shared by Jason Cyster^114^. CD19^−/−^ mice were crossed to MD4 mice. All lines were fully backcrossed to the C57BL/6J genetic background for at least 6 generations, used for experiments at 6-9 weeks of age, and housed in a specific pathogen-free facility at University of California, San Francisco, according to the University Animal Care Committee and National Institutes of Health (NIH) guidelines.

### Enzyme-Linked Immunosorbent Assay (ELISA)

Blood serum was tested by ELISA in order to quantitate Ag-specific IgG responses to various immunizations. 96-well plates (Nunc MaxiSorp, Invitrogen) were coated overnight at 4°C with either 200 ng of sHEL or 320 ng of sRBD per well in PBS. After blocking with 1% Bovine Serum Albumin (BSA, Fisher) in PBS, mouse sera of specified dilution factors were added to each well for incubation at 22°C for 2 hours. After three washes using PBS with 0.05% Tween 20, secondary goat anti-mouse IgG Fc-HRP antibody (# 1033-05, Southern Biotech) was added in blocking buffer at 1:6000 dilution and incubated for 1 hour at 22°C. Following three washes, 100 µL of substrate 3,3’,5,5’-Tetramethylbenzidine (Thermo Scientific) was added to each well and incubated in the dark for 10 minutes. The reaction was stopped by addition of 100 µL 2M sulfuric acid in each well. The optical density of each well at 450 nm was measured using a microplate reader (Bio-Tek Synergy HT). All the OD values reported were background subtracted by comparison between two wells that were coated with soluble protein and PBS, respectively. To estimate Ag-specific IgG concentration in mouse sera based on ELISA standard curves (Extended Data Fig. 2), all sera samples were properly diluted to reach final OD values between 0.2 and 0.5 before interpolation.

To determine the titers of Ag-specific IgG, we used a serial dilution of serum (from 1:100 dilution to 1:10,000,000 by a dilution factor of 10) for ELISA with the sHEL or sRBD protein. Cutoff values were calculated using the following equation as reported ^115^: Cutoff = X̄ + SD · *f*, where X□ and SD are the mean and standard deviation of control well OD reading values, *f* is the standard deviation multiplier corresponding to different confidence levels. Specifically in our assays, *f* = 2.631 when the number of control wells was 4 and the confidence level was 95%. The titer value was determined as the highest dilution factor of the serum that still yielded an OD450 value higher than the above cutoff value in ELISA.

To determine the subclasses of IgG antibodies elicited in mice upon immunization by various agents, we used the following secondary antibodies during ELISA: goat anti-mouse IgG1 HRP antibody (#1071-05, SouthernBiotech), goat anti-mouse IgG2a HRP antibody (#1081-05, SouthernBiotech), goat anti-mouse IgG2b HRP antibody (#1091-05, SouthernBiotech), goat anti-mouse IgG2c HRP antibody (#1078-05, SouthernBiotech), and goat anti-mouse IgG3 HRP antibody (#1101-05, SouthernBiotech).

### Preparation of HIV-1 virions pseudotyped with SARS-CoV-2 envelope

The HIV-1 pseudotyped with SARS-CoV-2 envelope was prepared following published protocols ^116^ but with modifications. Briefly, HEK 293T/17 cells (ATCC, Manassas, VA) were cultured at 37°C with 5% CO_2_ in DMEM supplemented with 10% FBS (HyClone Laboratories, Logan, UT). 10^6^ 293T cells in a 2-mL culture volume were seeded overnight in a 35-mm dish before transfection using the TransIT LT-1 transfection reagent (Mirus Bio, Madison, WI). For each dish, 2 μg of the provirus-containing plasmid pNL4-3 luc R^-^E^-117^ (ARP-3418, NIH AIDS Research and Reference Reagent Program) was used to make the transfection reagent mixture, together with 1 μg of envelope expression plasmid pcDNA3.1 SARS-CoV-2 S D614G ^116^. The plasmid pcDNA3.1 SARS-CoV-2 S D614G was a gift from Jeremy Luban (Addgene plasmid # 158075; http://n2t.net/addgene:158075; RRID:Addgene_158075). The RBD amino acid sequence 328-537 encoded in this plasmid is identical to the sequence of RBD that we studied here. The transfection reagent mixture was incubated at room temperature for 15 min before drop wise addition to the culture medium, as we did previously ^118^. At 6 hours post transfection, the culture medium together with the transfection reagents was replaced with fresh complete medium and the incubation was continued at 37°C with 5% CO_2_. At 48 hours post transfection, the entire culture medium containing single-cycle HIV-1 viruses was collected and filtered through a 0.45-µm syringe filter (Millex-HV PVDF, Millipore). The filtrate was then aliquoted on ice, flash-frozen in liquid nitrogen and stored in a -80°C freezer. The concentration of virion particles was quantitated using a HIV-1 p24 ELISA kit (CAT#XB-1000, XpressBio) as we described previously ^118^.

### Virion neutralization assay

The virion neutralization assay follows the protocols we established previously but with important modifications ^118^. HIV-1 virions pseudotyped with SARS-CoV-2 envelope containing 9 ng of HIV-1 p24 were incubated with various mouse sera at 37°C for one hour, and then diluted with complete medium by 8-fold in volume for the sera to initiate infection of Huh-7.5 cells at 37°C for 2 hours. At the end of 2 hours, fresh medium was added to each well in a 24-well plate and the incubation was continued at 37°C with 5% CO_2_. Luciferase activity was measured 48 hours after infection. Briefly, the culture medium was removed and replaced with 100 μL of complete medium. 100 μL of Bright-Glo^TM^ reagent (CAT#E2610, Promega) that was just warmed to room temperature was then added to each well. The cells were incubated for three minutes at room temperature to allow cell lysis. After three minutes, 100 μL of lysate from each well was transferred to a single well in a 96-well black microtiter plate (Costar). Luminescence was measured using a Synergy^TM^ HT multi-mode plate reader (BioTek Instruments Inc., Vermont) and background luminescence was subtracted using Huh-7.5 cells without virus infection. For comparison among different immunization groups in Fig. 2c and Fig. 4e, the luminescence readings for cells incubated with sera from PBS-injected mice was set as 100%. The luminescence readings from other groups were all normalized based on this and plotted as a percentage of remaining infectivity. For comparison among different time points post immunization, the luminescence readings for cells incubated with sera from each group of mice before injections was set as 100%. The luminescence readings from other time points for the same group were all normalized based on this and plotted as a percentage of remaining infectivity.

### In Vitro B Cell Culture and Stimulation

Splenocytes and/or lymphocytes were harvested into single cell suspension through a 40µm cell strainer, subjected to red cell lysis using ACK (ammonium chloride potassium) buffer in the case of splenocytes, and plated in round bottom 96 well plates in complete RPMI media with stimuli for various time as indicated. Complete RPMI media were prepared with RPMI-1640 + L-glutamine (Corning-Gibco), Penicillin Streptomycin L-glutamine (Life Technologies), 10 mM HEPES buffer (Life Technologies), 55 mM b-mercaptoethanol (Gibco), I mM sodium pyruvate [ (Life Technologies), Non-essential Amino acids (Life Technologies), 10% heat inactivated FBS (Omega Scientific).

For intracellular staining to detect pERK, stimulations for these assays were performed with 2 × 10^6^ splenocytes isolated as above per 100µL final volume. Following *in vitro* stimulation for 20 min, 1-2 x 10^6^ cells / well were resuspended in 96-well plates, fixed in 2% PFA for 10 min, washed in FACS buffer, and permeabilized with ice-cold 90% methanol at −20 °C overnight, or for at least 30 minutes on ice. Cells were then washed in FACS buffer, stained with intracellular Ab for 40 minutes, washed in FACS buffer, and then stained with Allophycocyanin (APC) AffiniPure F(ab’)C Fragment Donkey Anti-Rabbit IgG (H+L) (Jackson Immunoresearch) along with directly conjugated Abs to B220 in order to detect surface lineage and/or subset markers for 40 minutes. Samples were washed and then refixed with 2% PFA for 15 min. Abs to B220 (RA3-6B2) were conjugated to APC or PE. CD69 (H1.2F3) were conjugated to PE-Cy7 (Tonbo). Abs for intracellular staining: pErk1/2 T202/Y204 mAb (clone 194G2, cat. 4377S). The stimulatory Ab goat anti-mouse IgM F(ab’)_2_ was from Jackson Immunoresearch. After staining, cells were analyzed on a Fortessa (Becton Dickinson). Data analysis was performed using FlowJo (v10) software (Treestar Inc.).

### Statistical analysis

Statistical analysis was carried out using the Statistics toolbox in MATLAB (Mathworks, MA). A comparison of two independent variables was carried out using a two-sample T-test. Data sets with more than two independent variables were analyzed using a one-way analysis of variance as indicated. P-values less than 0.05 were considered statistically significant. Throughout, all errors in the main text were reported as standard errors unless otherwise noted; all data points in figures were reported as mean ± standard error unless otherwise noted.

## Acknowledgements

We thank Prof. Irina Grigorova for critical reading of the work during the early development of the manuscript. We thank Prof. Marilia Cascalho for very helpful discussion regarding the data and this work. We thank Dr. Charlie Rice at Rockefeller University for the kind gift of Huh-7.5 cells. The following reagent was obtained through the AIDS Research and Reference Reagent Program, Division of AIDS, National Institute of Allergy and Infectious Diseases (NIAID), National Institutes of Health (NIH): pNL4-3.Luc.R-E-from Dr. Nathaniel Landau. The plasmid pcDNA3.1 SARS-CoV-2 S D614G was a gift from Dr. Jeremy Luban through Addgene.

## Author contributions

Conceptualization: WC

Methodology: WW, ARM, SY, JLM, RM, BC, WC

Investigation: WW, ARM, SY, JLM, RM, JZ, WC

Visualization: WW, ARM, WC

Funding acquisition: WC, JZ

Project administration:WC

Supervision: WC

Writing: WW, ARM, BC, JZ, WC

## Data and materials availability

All data are available in the main text or the supplementary materials.

## Additional Materials for

**Extended Data Fig. 1.**
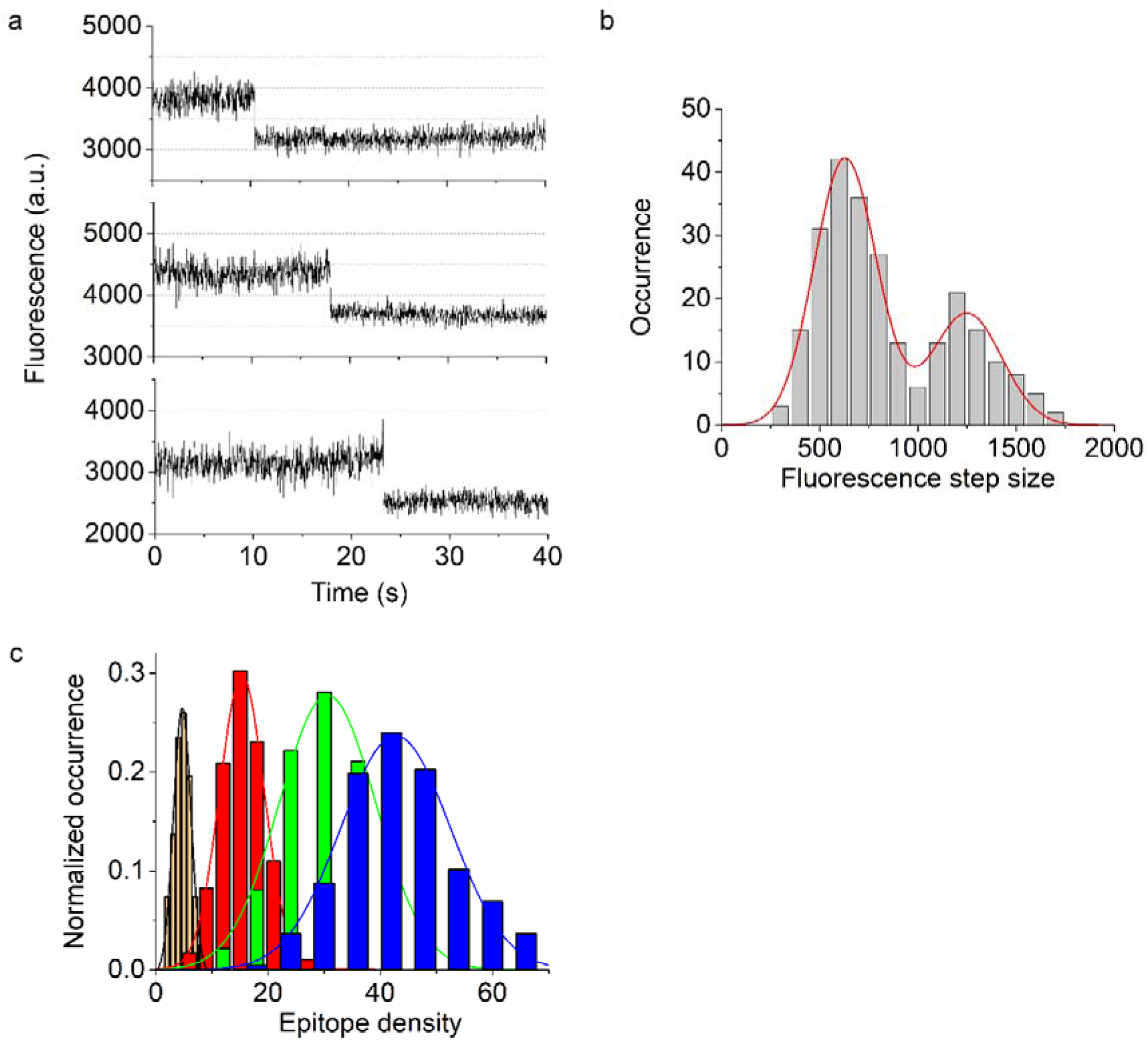
Characterization of RBD-conjugated SVLS using a single-molecule fluorescence approach. (**a**) Representative single-molecule fluorescence trajectories from Alexa-594 labeled RBD-specific Fab (Alexa-594-Fab). We prepared Alexa-594-Fab using a monoclonal IgG2a specific for SARS-CoV-2 RBD (BioLegend #944803, mAb1 herein). This high-affinity Fab allowed us to quantitate epitope density (ED) on individual SVLS conjugated with RBD, using single-molecule fluorescence techniques that we established previously ^70–72^. Briefly, the Fab was prepared from the IgG form of the antibody using a Pierce^TM^ Fab Micro preparation kit (ThermoFisher Scientific, CAT#44685) following manufacturer instructions. The purified Fab was then labeled using Alexa Fluor 594 succinimidyl NHS ester (ThermoFisher Scientific, CAT#A37572) following the protocol that we established previously ^71^. The labelling ratio between the dye and protein was determined to be 3.3 dye molecules per Fab based on single-molecule two-photon fluorescence measurement as we described previously ^71^. As shown in Fig. 1e in the main text, we deposited pRBD(DNA1)4 bound with Alexa-594-Fab on poly-L-lysine coated coverslips and used epi-fluorescence imaging to directly visualize individual SVLS through excitation of the Alexa-594 fluorophore by a 592-nm laser. Individual SVLS showed up as bright spots in a dark background. Neither SVLS alone nor free Alexa-594-Fab alone at the same concentrations yielded these bright spots under the microscope. Moreover, no fluorescent spots were visible when Alexa-594-Fab was incubated with control SVLS without RBD conjugation, confirming that these fluorescent spots were specific for SVLS conjugated with RBD but not protein aggregates. In order to determine the number of RBD molecules per SVLS, we chose to work at a laser power of 100 mW. This illumination condition allowed us to collect the initial fluorescence intensity from individual spots, and the constant illumination of the sample under this condition also allowed us to observe the photobleaching of individual fluorophores with time. As shown in Extended Data Fig. 1a, these photobleaching events display a hallmark feature of ‘steps’, where the fluorescence persists for a finite time followed by a sudden decrease in fluorescence intensity. These steps correspond to the photobleaching of individual molecules. We have used a step-detection algorithm that we developed previously ^119^ to identify steps from these real-time fluorescence traces. Measurement of this ‘step’ size yields the fluorescence intensity of a single Alexa-594 fluorophore, which can be used as an internal reference to convert the initial fluorescence intensity of individual liposomes to the number of Alexa-594 fluorophores, as we have done previously for individual HIV-1 virions ^72^ and individual protein-conjugated liposomes ^70^ using the same quantitation methodology. (**b**) The histogram of the individual photobleaching step sizes for Alexa-594 molecules (N=247), which could be well described by the sum of two Gaussians (red curve), one centered at 630 ± 15 analog-to-digital units (a.u., mean ± standard error) and the other centered at 1249 ± 40 a.u. Under a statistical significance level of 5%, Pearson’s Chi-square test ^120^ selected the double-Gaussian distribution (P-value=0.88) and rejected the single-Gaussian distribution (P-value=0.02) as the model to describe the data. The difference in peak values is close to twofold. As we have observed previously, the secondary peak may result from the photobleaching of two Alexa-594 molecules that occurred almost simultaneously, which could not be resolved by either the finite camera exposure time or the step-finding algorithm ^71,121^. The fluorescence intensity of a single Alexa-594 fluorophore can then be used to calculate the total number of Alexa-594 molecules per SVLS based on a ratio comparison ^122,123^ with the initial fluorescence intensity of Alexa-594 associated with individual SVLS. The number of RBD molecules per SVLS was then calculated as the total number of Alexa-594 molecules per SVLS normalized by the average number of Alexa-594 molecules per Fab. (**c**) The distributions of ED for pRBD(DNA1)1 (orange), pRBD(DNA1)2 (red), pRBD(DNA1)3 (green), and pRBD(DNA1)4 (blue). These distributions can be well described with Gaussian probability density function, with means and standard deviations of 5 ± 2 for pRBD(DNA1)1 (N=204), 15 ± 4 for pRBD(DNA1)2 (N=182), 31 ± 9 for pRBD(DNA1)3 (N=185), and 43 ± 10 for pRBD(DNA1)4 (N=217), respectively. These values are very close to the value of ED estimated using the ensemble approach as we established previously (Fig. 1f) ^59–61,70^.

**Extended Data Fig. 2.**
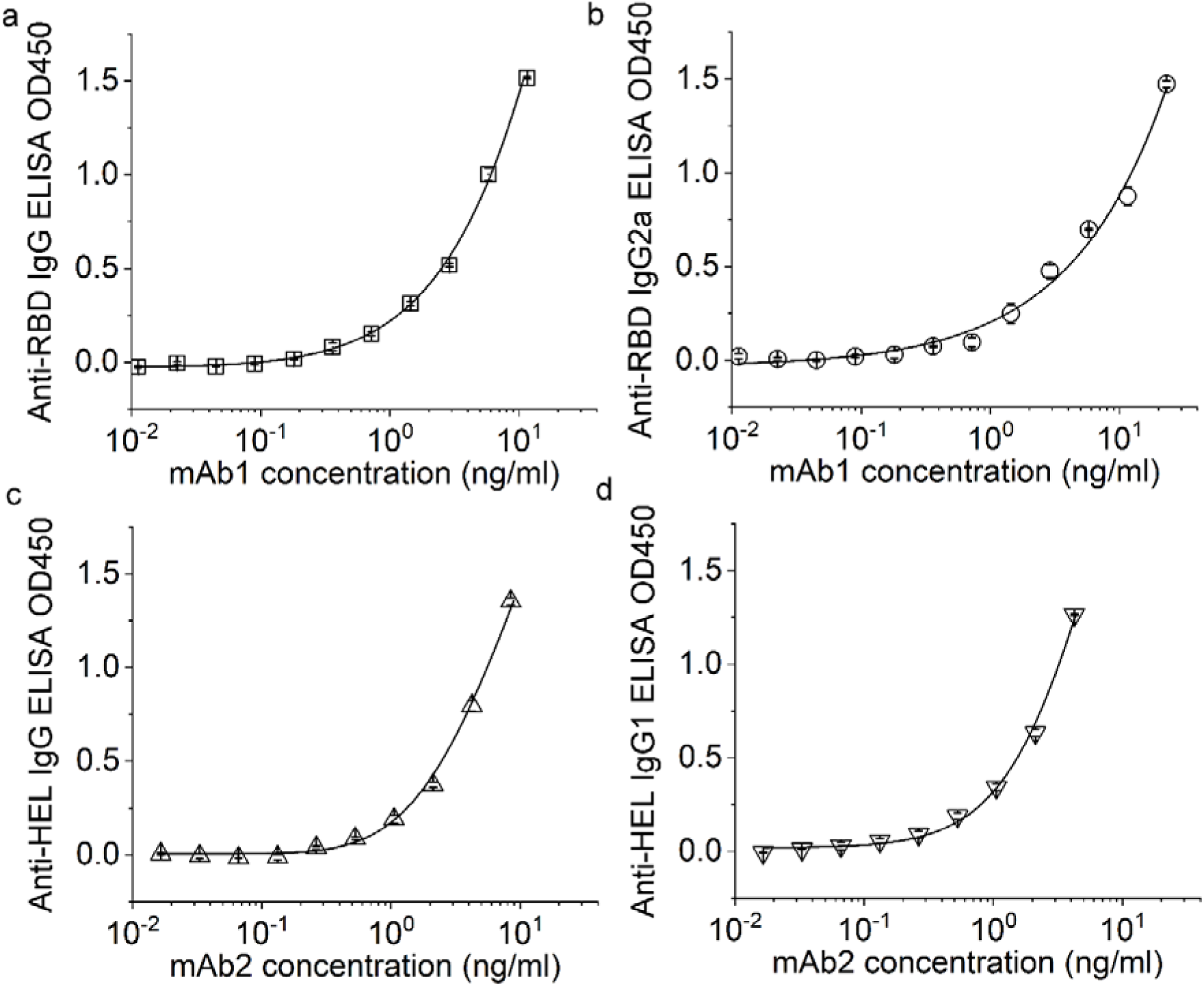
A representative set of standard curves for ELISA obtained using purified mouse monoclonal antibodies. (a) and (b) We used mAb1, the commercial RBD-specific monoclonal mouse IgG2a (BioLegend CAT#944803) as the known reference to construct standard curves for ELISA to indicate the sensitivity of current ELISA assays for detection of RBD-specific IgG and IgG2a. (c) and (d) We used mAb2, the HEL-specific monoclonal mouse IgG1 (clone HyHEL10, a special gift from Prof. Irina Grigorova) as the known reference to construct standard curves for ELISA to indicate the sensitivity of current ELISA assays for detection of HEL-specific IgG and IgG1. The results are representative of three or more independent repeats of the same experiments. The data were all fit to Richard’s five-parameter dose-response curves (black curves throughout Extended Data Fig. 2) in GraphPad Prism 9 for interpolation of Ag-specific IgG concentrations ^124^.

**Extended Data Fig. 3.**
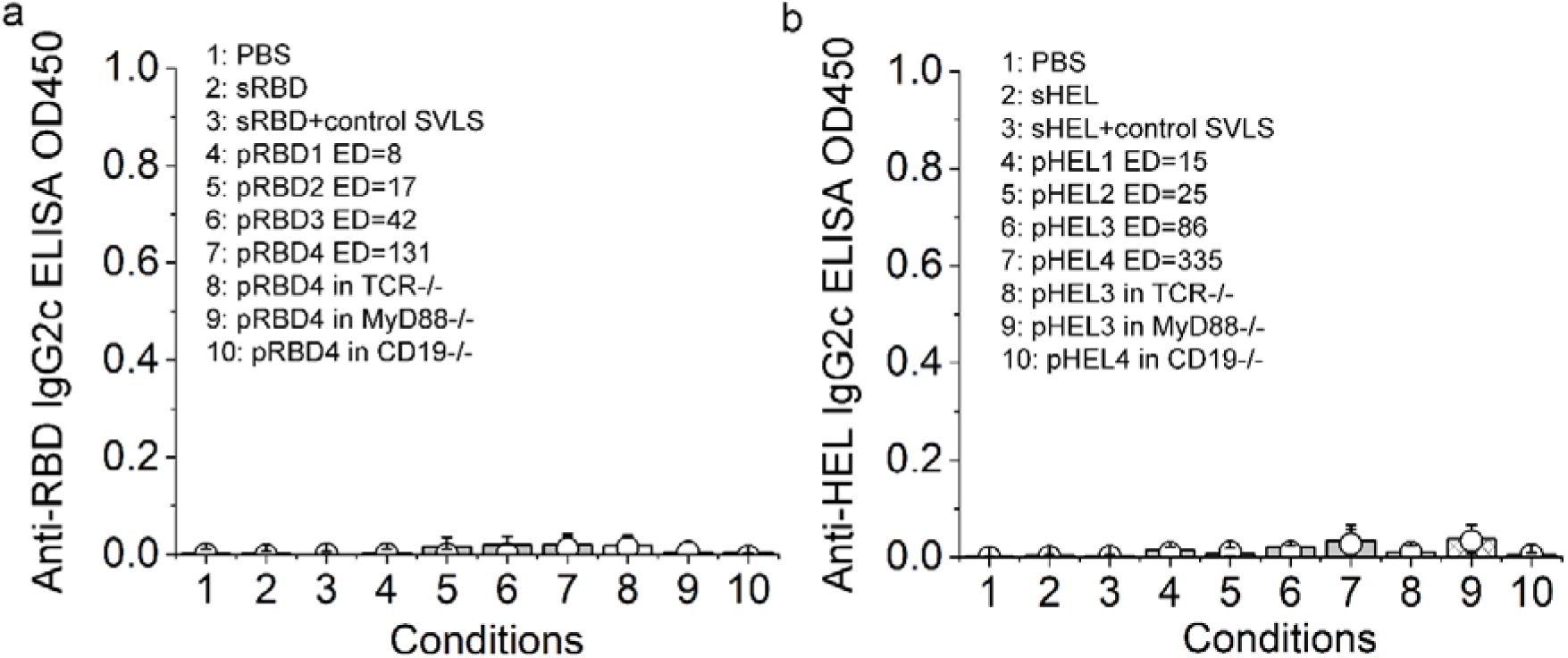
IgG2c was below the limit of detection in mice of B6 genetic background upon a single subcutaneous injection of SVLS without iNA. (**a**) ELISA OD values for RBD-specific IgG2c antibody from 1:100 diluted mouse sera after a single subcutaneous injection using various agents listed in the inset with no additional adjuvant. The dose of RBD was 0.24 µg per animal for Conditions 2 through 10. ED refers to epitope density. (**b**) ELISA OD values for HEL-specific IgG2c antibody from 1:100 diluted mouse sera after a single inoculation using various agents listed in the inset. The dose of HEL was 0.3 µg per animal for Conditions 2 through 10. Sera from Day 5 are shown in circles and those from Day 11 post immunization are shown in columns. Graphs depict N=4 mice and are representative of two independent experiments. Throughout Extended Data Fig. 3, Conditions 1 through 7 were for wtB6 mice. Conditions 8 through 10 were for gene knockout mice of B6 genetic background, with Condition 8 as TCR^-/-^, Condition 9 as MyD88^-/-^ and Condition 10 as CD19^-/-^, following the descriptions in the main text. The pairwise statistical difference between Condition 1 and others on Day 5 was determined by Student’s T-test and none of the conditions shows statistical significance, with p-value > 0.05. Error bars represent the standard errors (N=4).

**Extended Data Fig. 4.**
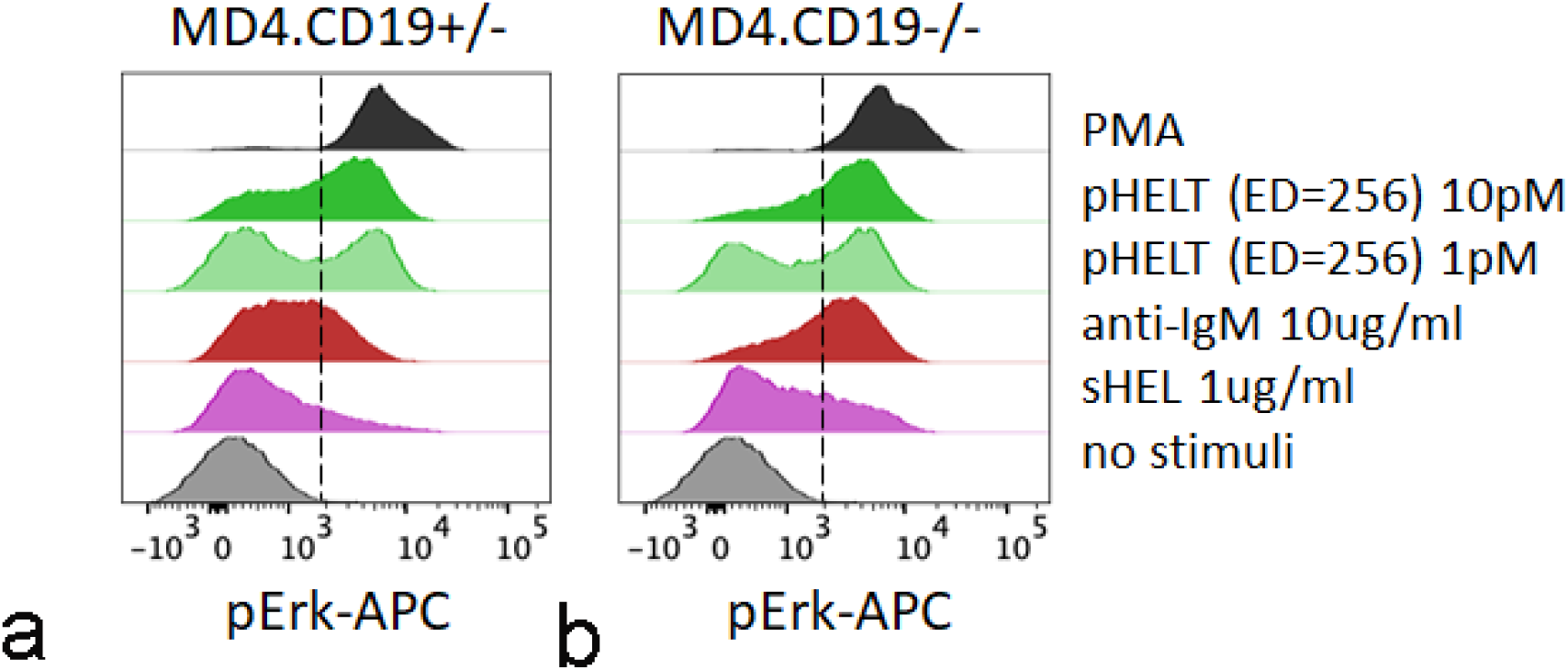
CD19 is dispensable for potent pHELT signaling *in vitro*. Splenocytes harvested from MD4 mice (harbouring Hy10 BCR inserted as Tg) either heterozygous or homozygous for CD19 deletion were incubated for 20 min with various stimuli as noted. Cells were then fixed, permeabilized, and intra-cellular pErk was measured by flow cytometry. Histograms depict pErk in B220^+^ cells from a single time course and is representative of 3 independent experiments.

**Table S1.** List of SVLS used in current and companion studies, together with the respective epitope density (ED), and the average number of internal nucleic acid (iNA) molecules per structure.

**S1a:**
| Name of SVLS | ED* | # of iNA per SVLS |
| --- | --- | --- |
| pRBD1 | 8±2 | 0 |
| pRBD2 | 17±2 | 0 |
| pRBD3 | 42±3 | 0 |
| pRBD4 | 131±9 | 0 |
| pRBD(DNA1)1 | 6±1 | 135±3 |
| pRBD(DNA1)2 | 18±2 | 140±4 |
| pRBD(DNA1)3 | 32±2 | 152±5 |
| pRBD(DNA1)4 | 46±2 | 139±3 |
| pRBD(DNA2) | 51±3 | 138±3 |
| pRBD(RNA1) | 55±2 | 147±4 |

**S1b:**
| Name of SVLS | ED* | # of iNA per SVLS |
| --- | --- | --- |
| pHEL1 | 15±2 | 0 |
| pHEL2 | 25±2 | 0 |
| pHEL3 | 86±4 | 0 |
| pHEL4 | 335±17 | 0 |
| pHEL(DNA1)1 | 5±1 | 118±3 |
| pHEL(DNA1)2 | 9±2 | 128±3 |
| pHEL(DNA1)3 | 20±2 | 136±4 |
| pHEL(DNA1)4 | 50±2 | 130±3 |
| pHEL(DNA2) | 47±2 | 126±4 |
| pHEL(RNA1) | 56±3 | 135±4 |

**(a)** RBD-conjugated SVLS and **(b)** HEL-conjugated SVLS. Throughout, the epitope density (ED) refers to the number of antigenic molecules specifically attached to the SVLS surface through maleimide chemistry. The # of iNA per SVLS refers to the average number of nucleic acid molecules encapsulated inside each SVLS. The uncertainties shown for both ED and iNA are standard errors determined from three independent repeats of the same ensemble experiment. Besides the above SVLS used in this study, control SVLS were SVLS that did not have any protein conjugation on surface nor have any iNA encapsulated inside SVLS.

**Table S2.** Statistical comparison of the titer values between SVLS(DNA1) and respective Ag-Qβ conjugates in wtB6 mice.

**S2a:**
|  | pRBD(DNA1)1 | pRBD(DNA1)2 | pRBD(DNA1)3 | pRBD(DNA1)4 |
| --- | --- | --- | --- | --- |
| Qβ-RBD, D5 | 0.356 | 0.537 | 0.024 | 1.06e-8 |
| Qβ-RBD, D11 | 2.88e-8 | 0.024 | 0.134 | N.A.* |

|  | pHEL(DNA1)1 | pHEL(DNA1)2 | pHEL(DNA1)3 | pHEL(DNA1)4 |
| --- | --- | --- | --- | --- |
| Qβ-HEL, D5 | 0.207 | 0.537 | 1 | 1 |
| Qβ-HEL, D11 | 0.134 | 0.356 | 0.356 | 0.356 |

**Table S3.** Statistical comparison of the titer values for SVLS(DNA1) of respective ED between wtB6 and BALB/c mice.

**S3a:**
|  | pRBD(DNA1)1 | pRBD(DNA1)2 | pRBD(DNA1)3 | pRBD(DNA1)4 |
| --- | --- | --- | --- | --- |
| BALB/c, D5 | 0.356 | 0.537 | 0.356 | 0.134 |
| BALB/c, D11 | 0.537 | 0.537 | 0.134 | 0.356 |

**S3b:**
|  | pHEL(DNA1)1 | pHEL(DNA1)2 | pHEL(DNA1)3 | pHEL(DNA1)4 |
| --- | --- | --- | --- | --- |
| BALB/c, D5 | 0.537 | 0.537 | 0.356 | 0.356 |
| BALB/c, D11 | 0.537 | 0.022 | 1.408e-8 | 0.022 |

**Table S4.** Statistical comparison of the IgG1 ELISA OD450 values for pRBD(DNA1) of respective ED between wtB6 and BALB/c mice.

|  | pRBD(DNA1)1 | pRBD(DNA1)2 | pRBD(DNA1)3 | pRBD(DNA1)4 |
| --- | --- | --- | --- | --- |
| BALB/c, D5 | 0.512 | 0.075 | 0.043 | 0.609 |
| BALB/c, D11 | 0.085 | 0.047 | 0.444 | 0.940 |

